# Novel hepatocellular carcinomas (HCC) Subtype-Specific Biomarkers

**DOI:** 10.1101/2023.08.28.555212

**Authors:** Jacob Croft, Odalys Quintanar, Liyuan Gao, Victor Sheng, Jun Zhang

## Abstract

**Introduction:** Hepatocellular carcinoma (HCC), the most common form of liver cancer, is a global health concern and a leading cause of cancer-related deaths. HCC accounts for a significant portion of liver cancers and has low survival rates of 5% to 30%, especially for HCC patients with a survival rate of 15%. Early detection is challenging due to the absence of symptoms in the early stages. The complexity and molecular diversity of HCC contribute to its poor prognosis. Understanding its molecular subtypes and mechanisms is crucial for improved management.

**Methods:** The study utilized publicly available data to investigate the potential diagnostic and prognostic biomarkers for hepatocellular carcinoma (HCC) based on their transcript per million (TPM) expression levels. A dataset of 407 HCC patient profiles was analyzed for survival trends and gene expression patterns.

**Results:** Through a comprehensive approach, over 900 potential prognostic candidates were identified. Further analysis narrowed down 647 prognostic and diagnostic candidate biomarkers. The study also explored the role of the CmPn signaling network in HCC, reaffirming that its components could act as prognostic markers. Additionally, the study-utilized machine learning to discover 102 transcription factors (TFs) associated with HCC, from candidate biomarkers.

**Conclusions:** The findings provide insights into the molecular basis of HCC and offer potential avenues for improved diagnosis, treatment, and patient outcomes. The expanded prognostic biomarker pool aids in pinpointing HCC-specific grading and staging biomarkers, facilitating targeted therapies for improved patient outcomes and survival rates

## Introduction

Hepatocellular carcinomas (known as hepatoma or HCC), the most common type of liver cancer, presents a worldwide health concern, ranking as the sixth most frequently diagnosed cancer and the second primary contributor to cancer-linked fatalities [1,2]. The escalation in liver cancer occurrences is mainly propelled by its principal subtypes, HCC, collectively responsible for 75% of all liver cancers [1,3,4]. With liver cancer’s 5-year survival rates spanning from 5% to 30%, the significance of early detection becomes critical for improving patient outcomes, especially considering individuals with HCC who experience survival rates of just 15% [5]. Identifying liver cancer in its early stages remains challenging due to its lack of symptoms until it reaches advanced phases [6–8]. HCC is particularly lethal due to its markedly low overall and long-term survival rates. The grim prognostic outcomes primarily stem from the extensive heterogeneity within HCC, which contributes to its diverse landscape. However, the understanding of molecular subtypes and the underlying mechanisms of HCC has fallen behind, hampering efforts to effectively combat this prevalent disease [9].

Biomarkers are crucial for predicting and monitoring liver cancer, aiding in timely detection, treatment response assessment, and disease progression tracking [10–12]. Swift intervention for HCC has shown to extend periods of recurrence-free health and enhance overall survival rates [13]. Nevertheless, HCC has been associated with resistance to chemotherapy, rendering it challenging to manage and resulting in discouraging prognoses [14]. Thus, the identification of sensitive and specific biomarkers could aid in the early discovery and control of liver cancer, ultimately ameliorating patient outcomes.

Currently, non-invasive diagnostics and prognostics for hepatic cancer, capable of distinguishing between distinct subtypes based on histology and morphology, are lacking. Alpha-fetoprotein (AFP) is the prevailing gold standard biomarker for liver cancer diagnosis, but its sensitivity and specificity vary, often requiring supplementary use [15–18]. The dearth of reliable hepatic cancer biomarkers underlines the urgency of identifying novel markers to enhance early detection and treatment selection for patients [11,12,15–17,19]. While AFP is a standard, its limitations in detecting various HCC grades and stages and accurately distinguishing primary liver cancer types are evident [15–17]. Furthermore, rarer subtypes of liver cancer encounter even fewer choices for early identification. Our recent discovery through differential expression pattern analysis in CmPn network components in two major types of liver cancers, HCC and Cholangiocarcinoma (CCA) tumor samples, at transcriptional and translational levels highlights their potential as innovative biomarkers for liver cancer, especially in HCC [20]. Consistent with our findings, the expression levels of various members of the CmPn network have been linked to the advancement of HCC from liver tumor grades to different cancer severities. Notably, progesterone receptor membrane component 1 (PGRMC1) and CCM3 have emerged as possible prognostic biomarkers for metastasis in HCCs [21–23]. Additionally, changes in the expression of all three CCM genes (CCM1-3) and PAQR7, members of CmPn as well, have been detected during the initial stages of liver cancer tumorigenesis [24]. These findings could lead to the development of more precise biomarkers, improving early diagnosis, treatment response assessment, and prognosis for liver cancer patients [20]. This study underscores the deficiency of sensitive and accurate biomarkers for anticipatory and diagnostic purposes in liver cancer, especially for its most common form, Hepatocellular carcinomas (HCC). This highlights the pressing need for fresh potential biomarkers capable of aiding early detection and non-invasive prognostic techniques, enabling the distinction among different liver cancer subtypes. By utilizing RNA-seq data, we identified genes exhibiting substantial differential expression at the transcriptional level in HCC tissues in comparison to normal tissues. While these genes hold potential as diagnostic biomarkers, their utility as prognostic biomarkers in distinguishing between HCC cancer subtypes is not strongly supported. With the advancement of scientific research, a variety of diagnostic strategies have emerged for handling various forms of cancer, which stand as some of the most lethal diseases affecting humans. These endeavors have played a role in the ongoing journey towards partially successful treatment of this life-threatening ailment [25–27]. Recognizing and cultivating biomarker discoveries represent crucial stages in preventing, promptly detecting, assessing treatments for, and predicting the regression and future outcomes of tumors [15,28,29]. The omics approach has emerged as a potent technique applicable to the discovery of biomarkers tailored to specific HCC subtypes [26,27,30]. Uncontrolled cellular proliferation in cancers, driven by disrupted cellular transcription regulation, necessitates targeting aberrant transcription factors (TFs) in distinct tumor cells for therapies and diagnostic/prognostic biomarkers [31–33]. Comprehensive omics analyses of transcription profiles have unveiled novel TF-derived prognostic biomarkers or panels across various cancers, including HCC [34–41]. Liver cancer, particularly HCCs, frequently emerges due to chronic factors such as cirrhosis triggered by hepatitis B or C, leading to its high incidence [25–29,42,43]. Viral-induced gene expression changes across DNA, RNA, and protein levels have been observed in HCC [44–51], emphasizing the significance of transcription factors (TFs) in liver cancer development. Therefore, a key aim of this study is to identify the TFs linked to potential diagnostic and prognostic biomarkers in HCC.

Public repositories containing HCC-related data have significantly expanded, encompassing comprehensive information such as epidemiology, clinical details, diagnostics, and Omics data [44–52]. The rapid expansion of HCC data repositories is anticipated to continue, continually enhancing the existing pool of diagnostic and prognostic biomarkers in the currently underserved domain of liver cancer research. Through bioinformatics analysis and the utilization of publicly available data, this study identifies potential biomarkers for major HCC types. Comparative analysis pinpointed candidate gene biomarkers associated with age and survival in HCC patients. Pathway analysis illuminated functions and disrupted pathways related to these biomarkers, providing insights into disease progression. Additionally, altered CCM protein levels in tumors propose their potential as biomarkers spanning multiple cancers [20,53–63]. Both progesterone receptor types (nPRs and mPRs) could potentially serve as biomarkers for genetically influenced cancer subtypes [20,61–65]. In this study, we employed publicly available data and our prior discoveries to perform bioinformatics analysis on combined public and our own datasets. This effort revealed 102 potential prognostic biomarkers, specifically transcription factors (TFs), for major HCC types. This expanded our understanding of molecular profiles and clinical significance in liver cancer, particularly for major HCC subtypes, from the initial pool of 647 HCC-specific diagnostic and prognostic biomarkers.

## Materials and Methods

### HCC patient data

Data pertaining to patients with hepatic cancer was sourced from the publicly available liver tissue data provided by The Cancer Genome Atlas (TCGA). A comprehensive collection of 407 HCC patient records was compiled, and a thorough examination of these specimens was conducted to establish their specific classifications in relation to the subtype of hepatic cancer. Our primary emphasis was placed on the identification of 407 patient transcriptome profiles that had received a diagnosis of the hepatic cancer subtype referred to as HCC. Although all these patient transcriptomes had been made accessible, the records did not contain information regarding the stage or grade of these cancers at the time of diagnosis. Consequently, our analytical approach was geared towards assessing the prognosis and diagnosis of HCC patients. From the accumulated patient samples, totaling 407 HCC patients, there are individuals who survived (n=175) as well as those who passed away (n=232).

### Transcriptomic Analysis of HCC Subtype Using TCGA Data

To understand the duration of the disease, survival curves were employed, which represented the probability of survival over time. Our investigation into patient transcriptomes persisted as we meticulously gathered data on RNA sequences encoding proteins. Protein-coding genes potentially serving as prognostic biomarkers were evaluated for their gene expression using transcripts per million (TPM), with consideration of the samples’ survival status. The data were sourced from liver cancer samples in the TCGA database, while control liver tissue samples were obtained from the GTEx Program database. Survival status of TCGA samples was determined, classifying them as “HCC.Alive” for surviving (S) patients and “HCC.Dead” for patients who had deceased (D). Healthy control samples were labeled as “Control.” The Kaplan-Meier survival plots for the outcomes of patients with hepatocellular carcinoma NOS who experienced fatal outcomes due to their condition were used as the second filter to categorize the patients. Roughly, a quarter (25%) of individuals who faced mortality from hepatocellular carcinoma survived beyond 1000 days. These data were vital in capturing alterations in functional units that were influenced by the progression of cancer. The survival outcome of each patient was categorized and used as a prognostic indicator. This allowed us to compare the gene expressions of over 19,000 genes, measured in transcripts per million (TPM), between those patients who survived and those who succumbed to the disease.

To establish potential prognostic biomarkers, a Welch’s t-test was utilized to determine the significance of differences in means between the two groups. The results of this analysis enabled us to pinpoint potential prognostic biomarkers linked to patient outcomes and the expression of tissue protein coding RNA sequences.

### Normal liver tissue controls for comparative analysis of prognostic biomarkers

Furthermore, apart from identifying the prognostic biomarkers, we procured control samples of liver tissue from the GTEx portal. This yielded a total of 272 control liver tissue samples. From these, we extracted the protein-coding mRNA species that had been identified as prognostic indicators. For the purpose of analysis, we chose statistically significant genes by making a comparative assessment between the patients with HCC and the expression profiles of control samples derived from healthy individuals.

### Pathway analysis for candidate prognostic biomarkers with Gene Expression data

To the total number of genes that displayed statistically significant differences between the survivor and deceased HCC patient cohorts. We performed gene set enrichment analyses (GSEA) on genes displaying the highest level of statistical significance. To gain a comprehensive understanding of the pathways impacted and contributing to distinct prognosis, we conducted the GSEA analysis across three distinct libraries (GO, KEGG, and DOSE). Continuing our inquiry, we implemented a filtering mechanism specifically targeting genes exhibiting unique expressions linked to HCC. This approach aimed to exclude genes that were shared between both cancer types. As a result, the genes that held significance in influencing patient prognostic outcomes were eliminated, as the primary filter. Following that, a secondary GSEA analysis on genes displaying the highest level of statistical significance was carried out, specifically aimed at identifying pathways that were uniquely affected by HCC, as the secondary filter.

### Statistical analysis

Through the application of the t-test, we investigated not only the significantly differential expressions between the healthy control and HCC cohorts but also delved into the significantly differential expressions within the survival and deceased HCC patient groups, subsequently comparing them to the control group. To analyze the differentia expression patterns of protein coding RNAs, we initially employed a Welch’s t-test to identify TPM values of mRNAs. This outcome facilitated the identification of differences in prognostic status, leading us to establish a filter for mRNAs displaying differential expression.

Subsequently, we exclusively compared the statistically significant differentially expressed mRNAs identified in the initial step, contrasting HCC survivors with the normal controls. This analysis was co-inducted to ascertain the significance of the differential expression in relation to the healthy control population. Our objective was to pinpoint exclusively distinct TPM values to establish diagnostic potential. In conclusion, we employed another Welch’s t-test on mRNAs showing significant differences between HCC survivors and controls. Next, we aimed to determine if these protein coding mRNAs exhibited significant differential expression in the deceased HCC cohort. This approach enabled the identification of unique mRNAs that showcased differential expression not only in survival and deceased HCC cases, but also in both cohorts when compared to controls. This established both prognostic and diagnostic capabilities.

### Transcription Factor prediction and Experimental Setup

The degree of difficulty for predicting transcription factors (TFs) stems from the intricate and resource-intensive process of experimental validation [66–68]. In this research, we utilized our developed ESM-TFpredict framework, designed to automate the identification of potential TFs [58,69]. To enhance training efficiency while working with limited resources, the sequence is divided into predefined windows. The amino acid representations within each window are averaged to create a concise overall representation. The prediction model then employs two layers of 1-D CNNs (with 128 and 256 filters respectively), each followed by a max pooling layer and a dropout layer (with a dropout rate of 0.5). Following these layers, a dense layer and another dropout layer are applied to further refine the extracted features and mitigate overfitting. Ultimately, the final prediction is generated using a sigmoid activation function.

### Dataset Preprocessing, experimental execution and evaluation

Our protein dataset was primarily sourced from UniProt (www.uniprot.org). To create a set of TF sequences with a high level of confidence, we thoroughly examined the Swiss-Prot dataset, concentrating on annotations that included the terms “transcription regulation” and “DNA binding” [70,71]. This careful curation resulted in a collection of 5,418 TF sequences, all of which possessed a flawless 5/5 annotation score. For sequences not related to transcription factors (NTF sequences), we intentionally filtered out instances containing the aforementioned keywords, resulting in a pool of 47,120 sequences. To achieve balance within the dataset, we randomly selected 5,564 NTF samples. Notably, over 98% of the sequences have a length of less than 3,000 amino acids (Suppl. Table 1). To maintain consistent analysis, we have capped the sequence length at 3,000 in our experiments, truncating longer sequences accordingly.

Experiments were conducted on a Linux server with an Intel(R) Core(TM) i9-10900X CPU running at 3.70GHz, augmented by a GeForce RTX 3080 GPU and 122GB of RAM. The ESM-TFpredict model was trained for 40 epochs using a batch size of 128 and a learning rate of 10^-4. The condensed representation was set at a fixed size of 300. Employing 5-fold cross-validation yielded average scores of F1 = 0.9531, specificity = 0.9537, sensitivity = 0.9525, and balanced accuracy = 0.9531. These results underscore the reliability and precision of our machine learning (ML)/deep learning (DL)-based TF prediction model.

## Results

### Age Distribution, Survival Trends, and Prognostic Possibilities within our HCC Patient

In the beginning stages of our exploration using the TCGA-derived dataset, we compiled a group consisting of 407 patient profiles affected by HCCs. Initially, our focus was on understanding the age distribution within this cohort and deciphering the patterns related to survival rates (Figure 1A). Subsequently, we investigated the lifespan of HCC patients, as depicted by the KM survival curves (Figure 1B). Our findings revealed a distinct trend: as time progressed, the overall prognosis associated with HCC diagnosis displayed a rapid decline (Fig. 1). Furthermore, we noticed that unlike other forms of liver cancer, there is a distinct and heightened vulnerability to HCC diagnosis between the sixth and eighth decades of life. As time passes, the likelihood of survival decreases; the graph illustrates that only a small number of HCC patients manage to survive beyond 3000 days.

**Figure 1.**
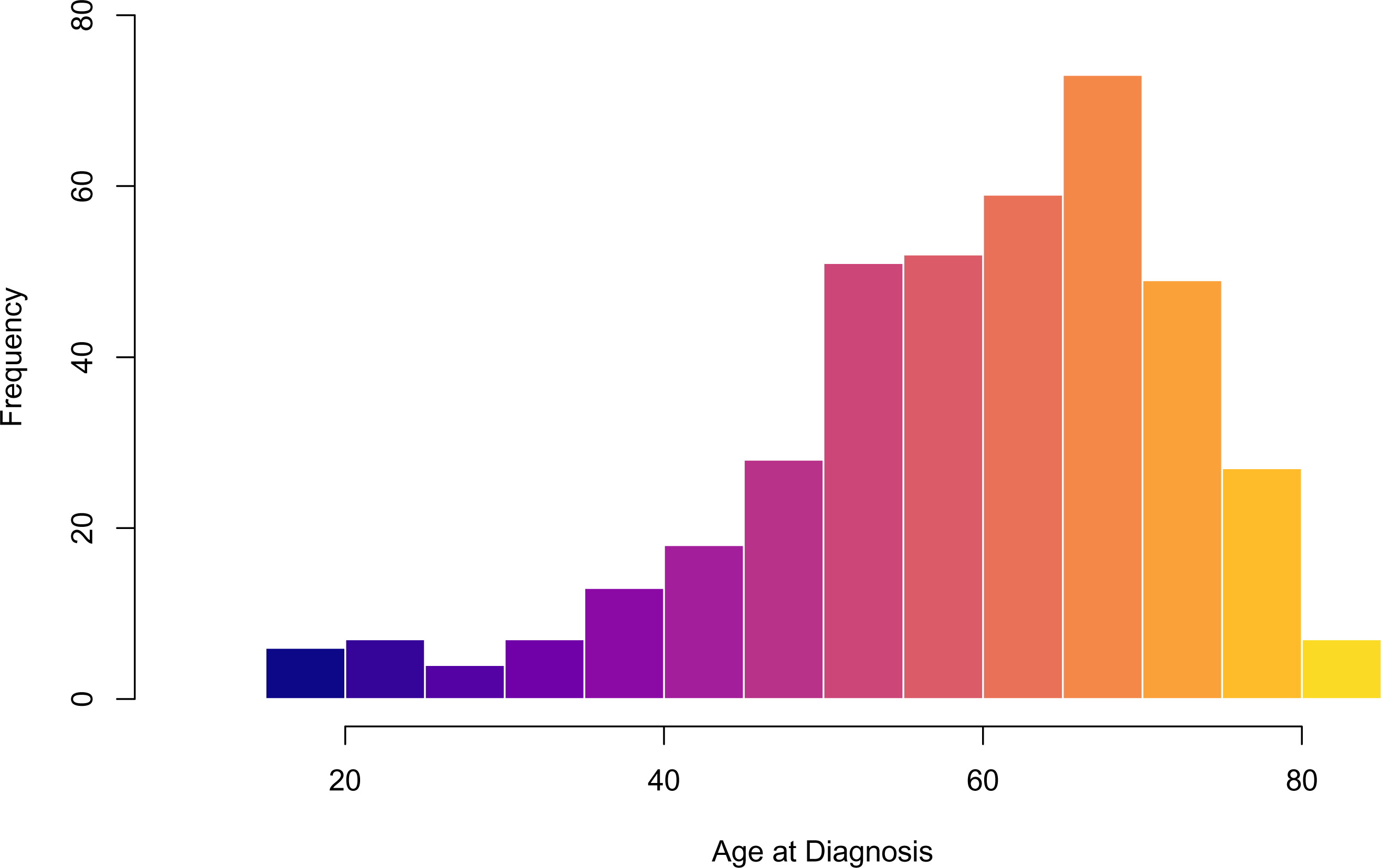
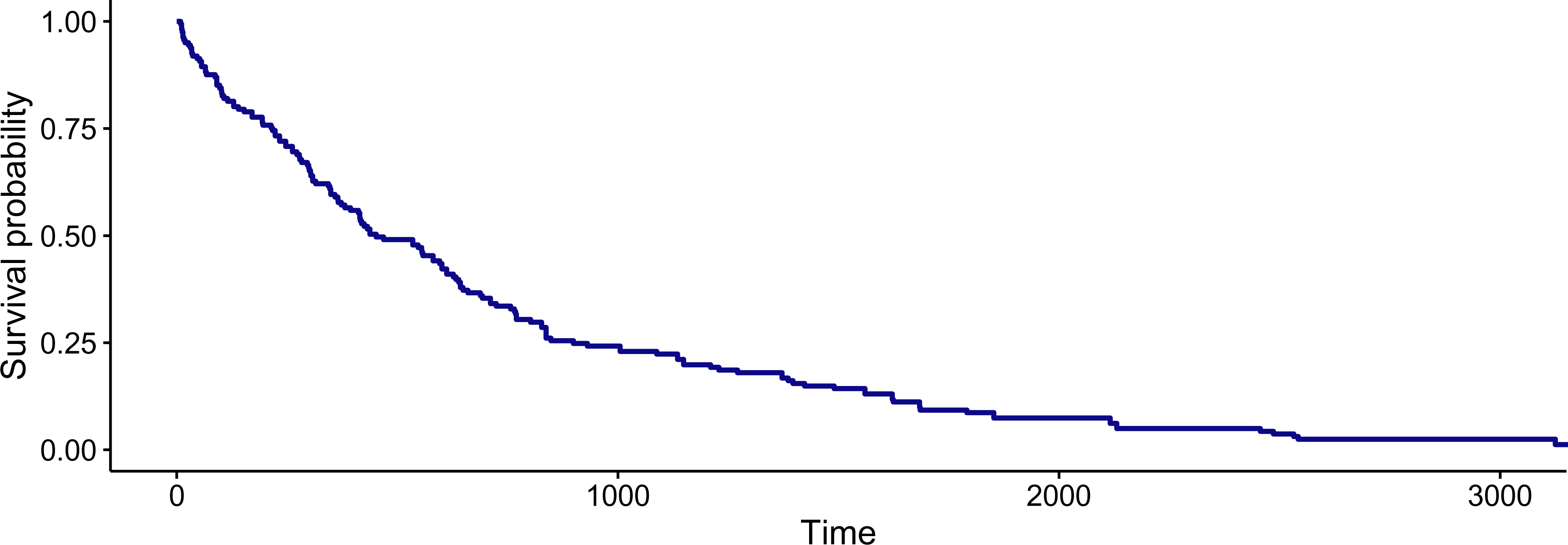
Demographic Analysis and Survival Patterns of Hepatocellular Carcinoma (HCC) Patients Based on TCGA Data. HCC represent the predominant form of liver cancer. The analysis utilized data obtained from data repertoire publically accessible patient transcriptomes, encompassing essential demographic details. While the specific stages and grades of the cancers were not determined. **A.** The age distribution among HCC patients is illustrated as identified at the time of diagnosis. Notably, the highest vulnerability to diagnosis was observed between the sixth and eighth decades of life. **B.** Kaplan-Meier Survival curve of HCC Patients displays the duration of survival in days, plotted on the x-axis, as a means of prognosticating the HCC patient’s condition over the course of the disease. As time progresses, survival likelihood decreases; the graph showed that few HCC patients survived over 3000 days.

### Identification of Prognostic and Diagnostic Biomarkers from Comprehensive Analysis of Protein-Coding RNA Sequences in HCC Patients cohort

Continuing our experiment, we thoroughly assessed upwards of 19,000 protein-coding mRNAs, unveiling a pool of over 900 potential candidates exhibiting notable prognostic significance. These candidates exhibited diverse transcript per million (TPM) expression patterns in relation to survival outcomes. We conducted a more in-depth analysis of these over 900 prognostic candidates, enhancing the results by assessing not only the statistical differentiation between survival HCC (S) patients and normal controls but also between deceased (D) HCC cases and normal controls. Within this pool of 900 candidates, we presented the top 12 to showcase their potential as the representative prognostic biomarkers (Fig. 2A). To conduct a more thorough investigation, our focus was directed towards 939 candidates displaying significantly differential expression. These candidates not only showcased statistically significant expression differences between normal controls and surviving HCC patients but also revealed same statistical significance in expression patterns between HCC patients who survived and those who passed away, we presented the top 12 to showcase their potential as the representative prognostic and diagnostic biomarkers (Figs. 2A-B and Supplementary Figures 1A-B). In a preceding study, we established that the fundamental elements in the CmPn signaling network, comprising *mPR*α *(PAQR5), mPR*β *(PAQR6), mPR*γ *(PAQR7), mPR*δ *(PAQR8), mPR*ε *(PAQR9),* and *CCMs (1-3)* genes are linked to HCCs. This connection provides an avenue for further investigation into their viability as potential biomarkers for HCC. As a potential group of positive control biomarkers, we subjected the essential components within the CmPn signaling network to re-evaluation, leveraging an expanded dataset and enhanced analytical tools. This re-evaluation confirmed their potential and underscored their roles as both prognostic and diagnostic biomarkers (Fig. 2C). Based on these observations, we deduced the potential for identifying biomarkers that are exclusive to prognosis, as well as biomarkers that possess both prognostic and diagnostic capabilities (Fig. 2 and Suppl. Fig. 1). In fact, this two-phase analysis brought to light a total of 647 strong candidates for prognosis and diagnosis (Suppl. Table 1).

**Figure 2.**
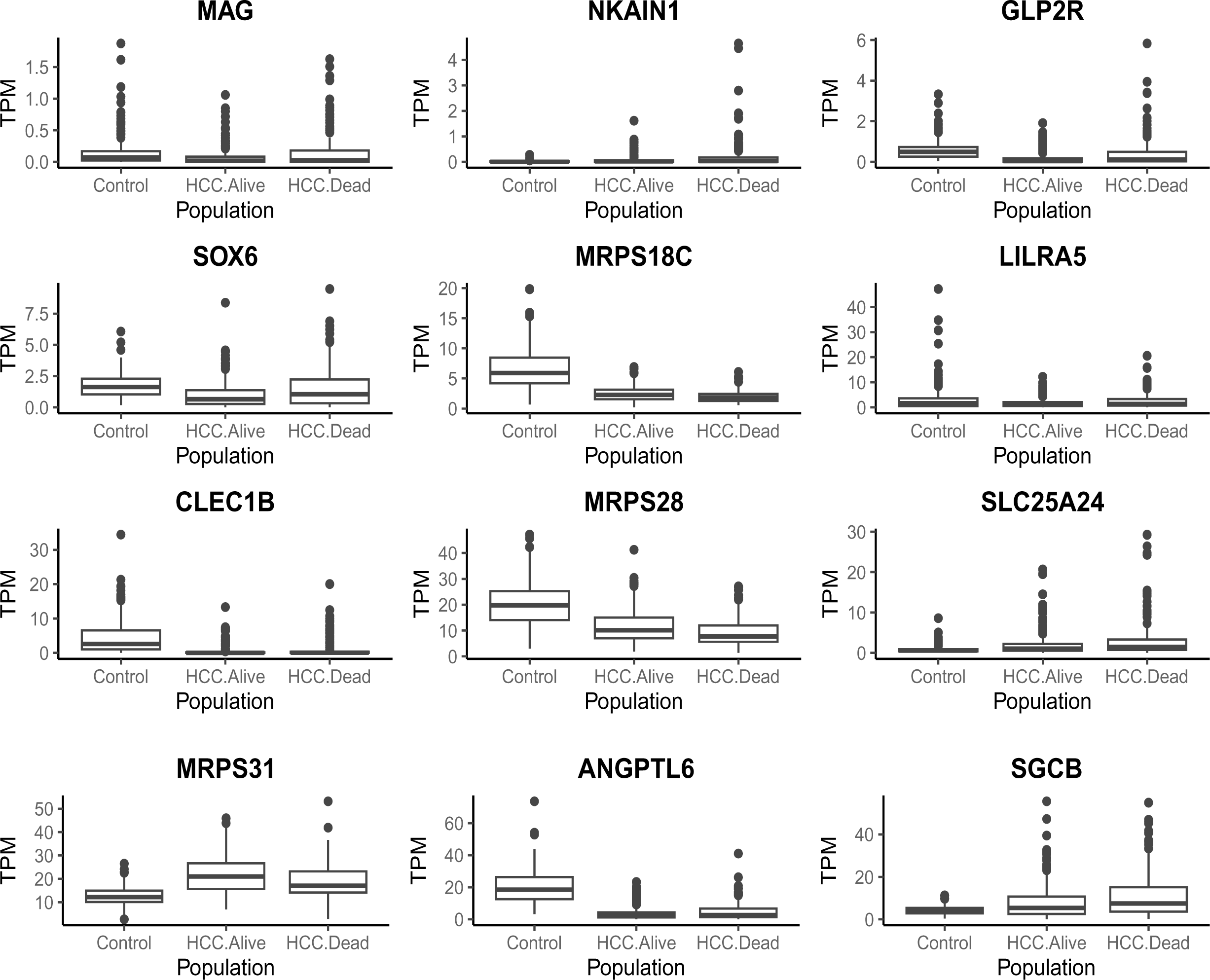
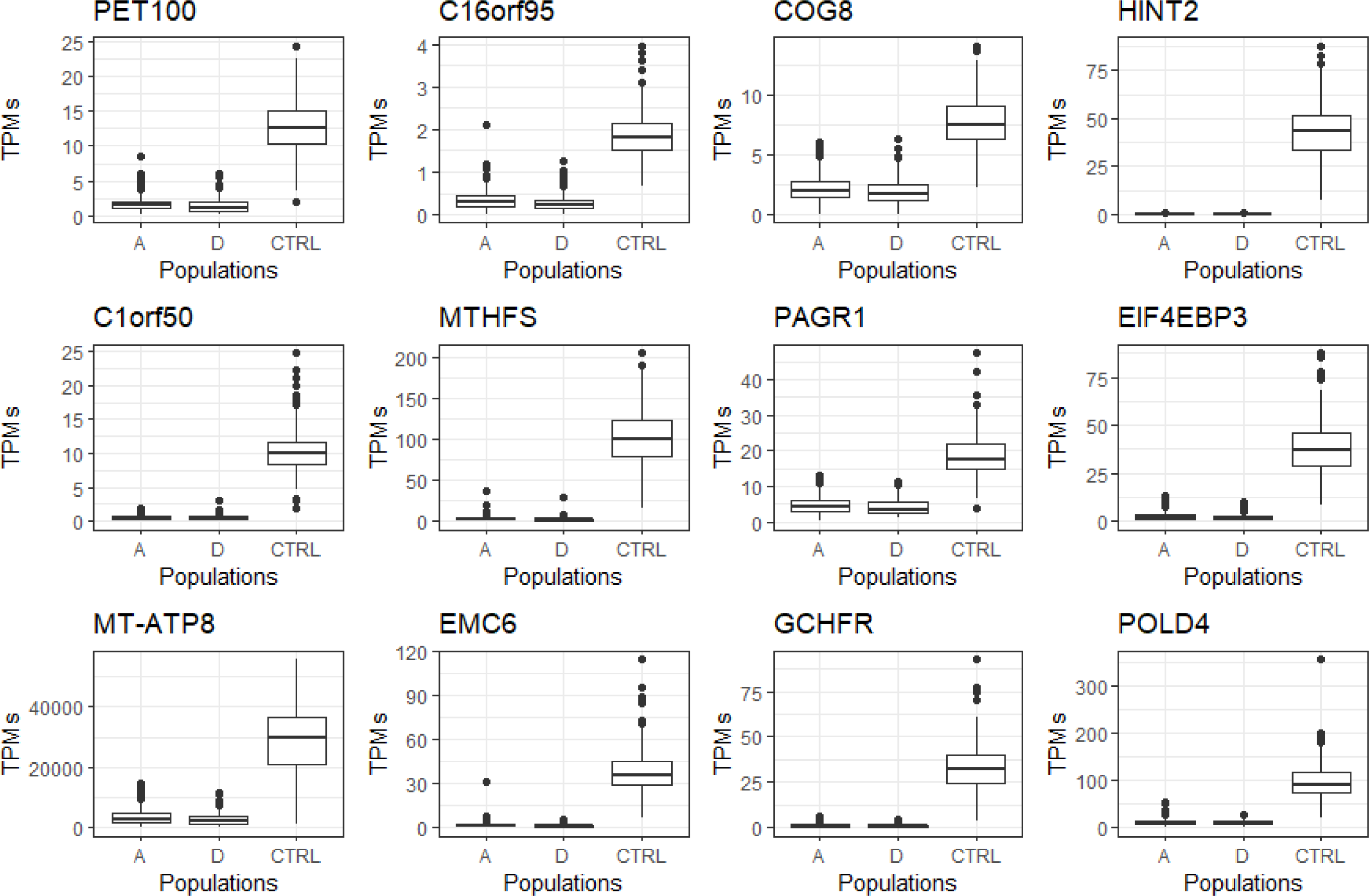
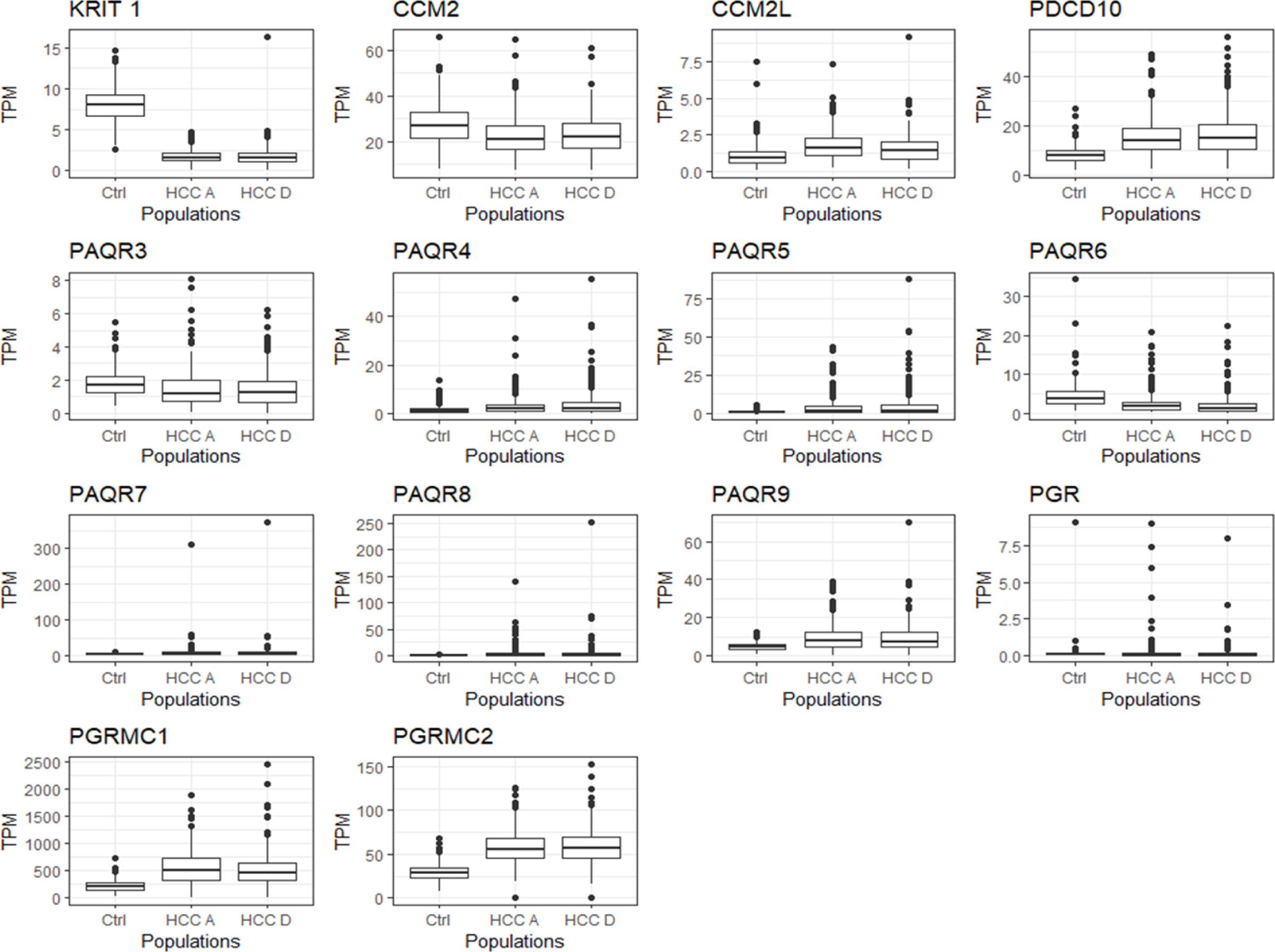
Assessing the Viability of Transcriptome-Based Prognostic Biomarkers in HCC Related to Protein-Coding Genes. Boxplots were utilized to illustrate the dispersion of data concerning protein-coding genes potentially functioning as prognostic biomarkers. Categorizing the tools into prognostic and prognostic/diagnostic groups enabled the identification of the highest number of detectable biomarkers. Genes’ expressions, measured as Transcripts per Million (TPM), were sorted based on the vital status of the samples. Our analytical approach involves both comprehensive and comparative examination of data sourced from the TCGA database, alongside the TPM values from samples of Healthy Controls (CTRL), surviving (S), and deceased patients (D) with HCC. Importantly, all the genes depicted in the graphs are viable contenders for prognostic biomarkers. **A.** The first experiment produced over 900 potential prognostic biomarkers exclusively for HCC liver cancer subtype. This determination was made by contrasting the transcriptome profiles of surviving (S) individuals with those of deceased (D) HCC cancer patients. The plot here illustrates the top 12 results out of top 25, while the remaining outcomes can be referenced in supplementary Figure 1A. **B.** The next experiment involved the comparison of over 900 candidate biomarkers between survivors and control groups, followed by a comparison between deceased and control populations. This analysis unveiled more than 650 potential prognostic and diagnostic biomarkers. The provided list includes the top 12 results out of 25, while the remaining outcomes can be referenced in supplementary Figure 1B. **C.** In our prior work, we identified members of the CmPn as potential biomarkers for determining the different stages of HCC. We sought to re-evaluate the validity of our earlier results and discovered that all 14 members of the CmPn pathway could serve as prognostic biomarkers for HCC. Furthermore, 9 out of these 14 members could function as both prognostic and diagnostic biomarkers. (Note: The remaining parts of these plots were illustrated in Supplemental Figures 2A-B).

### Discovery of Prognostic Biomarker through Pathway Enrichment in HCC

Next, we embarked on a series of comprehensive pathway enrichment analyses aimed at understanding the functional implications of genes with significantly differential expression as prognostic biomarkers unique to the HCC. This study involved employing Gene Set Enrichment Analyses (GSEAs) across three distinct libraries (GO, KEGG, and DOSE), specifically tailored to the genes recognized as unique prognostic biomarkers in HCC (Suppl. Figs. 2A-C). By conducting these analyses, our aim was to reveal the biological pathways and molecular functions linked to these genes through a combined comparative GSEA approach (Fig. 3A). Within this array of pathways, viral infections and inflammatory responses, hormonal reactions, and metabolic pathways exerted influence, unveiling their potential importance of identified prognostic biomarkers within the realm of HCC prognosis.

**Figure 3.**
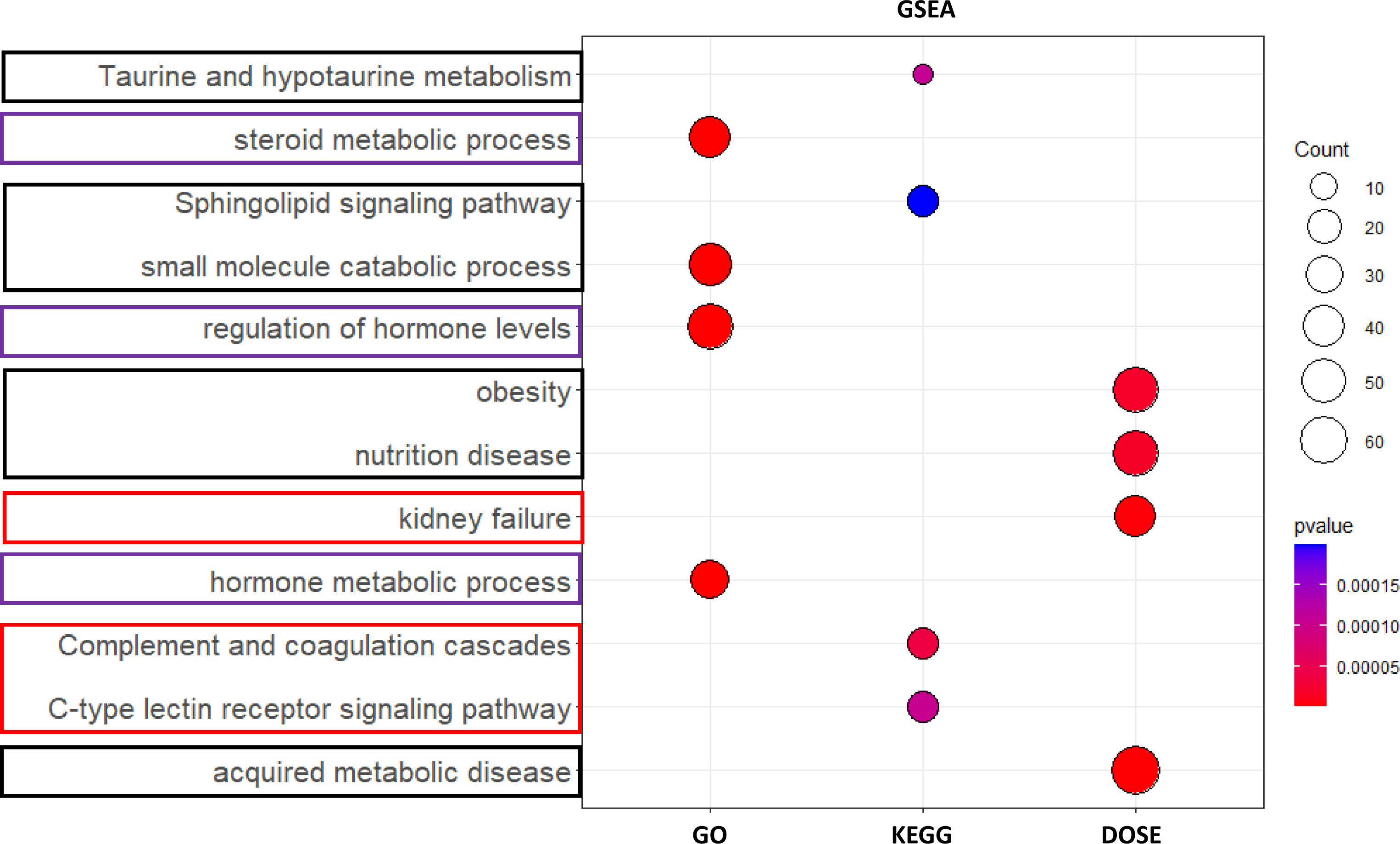
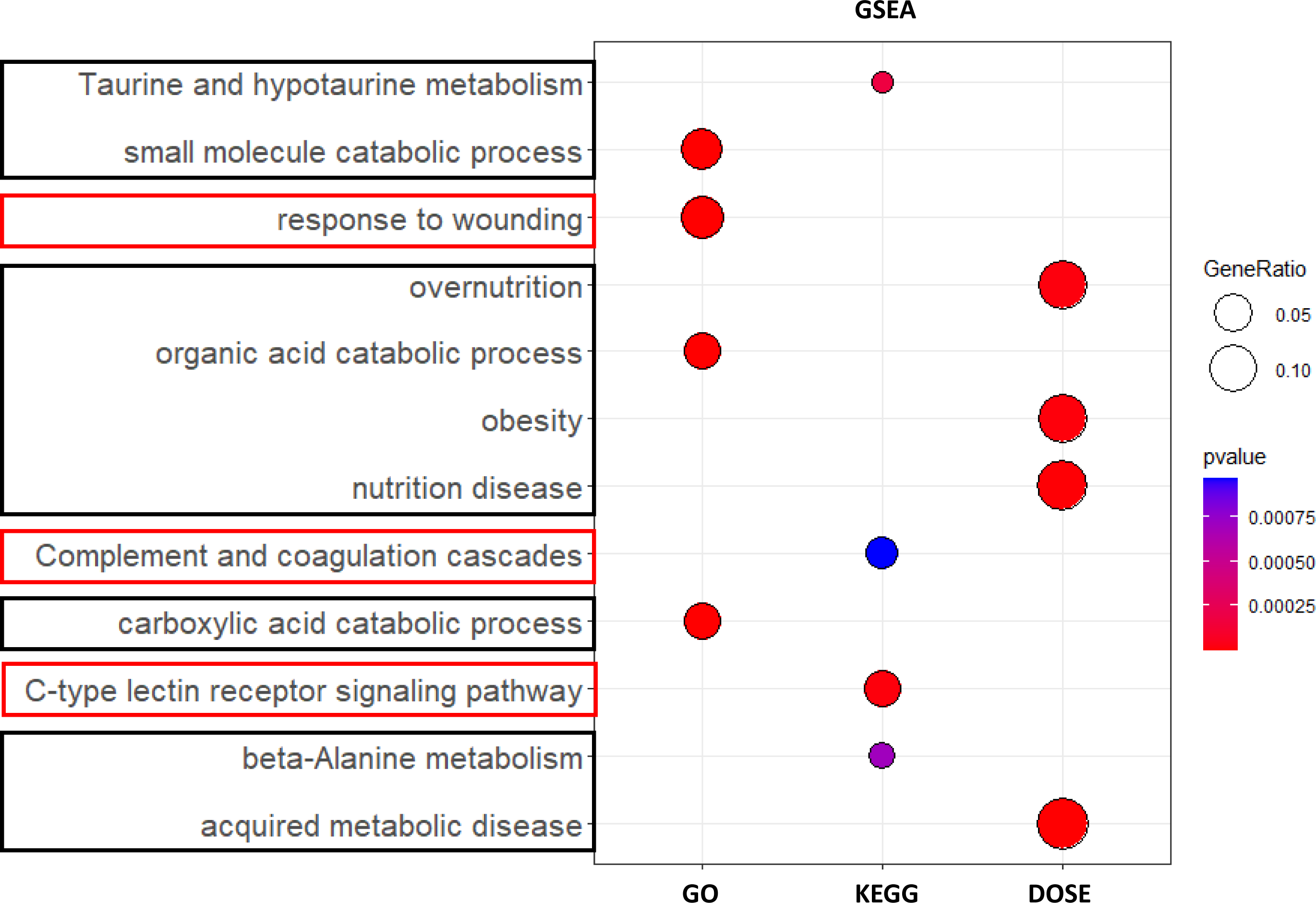
Uncovering unique HCC-subtype prognostic biomarkers through Gene Set Enrichment Analyses (GSEA). By utilizing three separate enrichment libraries, we highlight the pathways influenced by HCC tumorigenesis, with the intention of identifying the primary diseases associated with these genes. **A.** This plot visualizes the signal pathways linked to more than 900 identified mRNA species, which have been identified as potential prognostic biomarkers through a comparative expression analysis conducted between surviving (S) and deceased (D) HCC patients. **B.** Similarly, this plot depicts the signal pathways linked to more than 900 identified mRNA species, which have been recognized as potential diagnostic and prognostic biomarkers. This recognition is based on a comparative expression analysis conducted among survival (S) HCC patients and Healthy Controls (CTRL), as well as deceased (D) HCC patients and Healthy Controls (CTRL). Moving across from the left, we can see enriched analyses involving gene ontology (GO), Kyoto Encyclopedia of Genes and Genomes (KEGG), and Disease Ontology Semantic and Enrichment (DOSE). These analyses highlight pathways impacted by viral infections and inflammatory responses (indicated by red outlines), hormonal responses (purple outlines), and metabolic pathways (outlined in black). After refining the results of the earlier prognostic biomarker candidates, we identified that most of the affected pathways remain linked to metabolic processes and reaction in the context of wound-related effects (inflammatory responses), which corresponds with pathways influenced in other forms of cancer. In sum, the pathways that overlap between these two analyses offer promise for identifying diagnostic and prognostic biomarkers, which can be subjected to further validation analysis in the future.

### Identification of Dual diagnostic and prognostic biomarkers and Associated Pathways in HCC via Comprehensive GSEA

We then proceeded to perform an enrichment analysis aimed at identifying the signal pathways that were most profoundly disrupted by the HCC. This analysis entailed a meticulous comparison between pathways deemed normal and those exhibiting significant statistical differences within both surviving and deceased HCC patient groups. Additionally, we thoroughly examined the statistical significance of gene expression patterns across normal controls and distinct groups of surviving and deceased HCC patients. This investigation encompassed pathway enrichments spanning three distinct libraries (GO, KEGG, and DOSE) to define HCC-specific diagnostic and prognostic biomarkers (Suppl. Figs. 3A-C).

By conducting these analyses, our aim was to reveal the biological pathways and molecular functions linked to these genes through a combined comparative GSEA approach (Fig. 3B). From this array of pathways subjected to a more rigorous selection process, it became evident that only viral infections and inflammatory responses, as well as metabolic pathways, demonstrated their influence. Ultimately, the comprehensive outcomes identified the significantly impacted signaling pathways in HCC tumorigenesis. This process also revealed intricate molecular alterations associated with HCC’s development and progression (Fig. 3B). This highlights the potential importance of the identified prognostic and diagnostic biomarkers in monitoring the development and advancement of HCC.

### Some diagnostic/prognostic biomarkers act as TFs to regulate Associated Pathways in HCC

Using our devised ML/DL-based ESM-TFpredict model [58,69], we have effectively identified 102 transcription factors (TFs) out of the 647 HCC-specific diagnostic and prognostic biomarkers that we previously uncovered (Table 1, Supplementary Table 1). Our findings align with transcription profiling data observed in major cancer types, especially HCC [34,36,38,39,41,72]. Acknowledging the significant influence of transcription factors (TFs) in liver cancer progression [43,51,73], we conducted enrichment analyses to reveal the signaling pathways where the identified TFs play a role within the context of HCC. Through the utilization of three enrichment libraries (GO, KEGG, DOSE), our analyses were aimed at identifying 102 HCC-specific TFs that hold potential as prognostic biomarkers (Table 1, Suppl. Figs. 4A-C). The effort to elucidate perturbed signaling pathways in HCC linked to these TFs has been effectively accomplished through a consolidated GSEA evaluation (Table 1, Fig. 4). Interestingly, the four principal signaling pathways – spanning tumorigenesis (outlined in light blue), metabolic pathways (black), inflammatory responses (red), and hormonal responses (purple) – show a convergence with pathways previously identified within diagnostic and prognostic biomarkers (Fig. 3). This alignment strongly suggests that these pathways are governed by the identified TF biomarkers in the context of HCC tumorigenesis.

**Figure 4.**
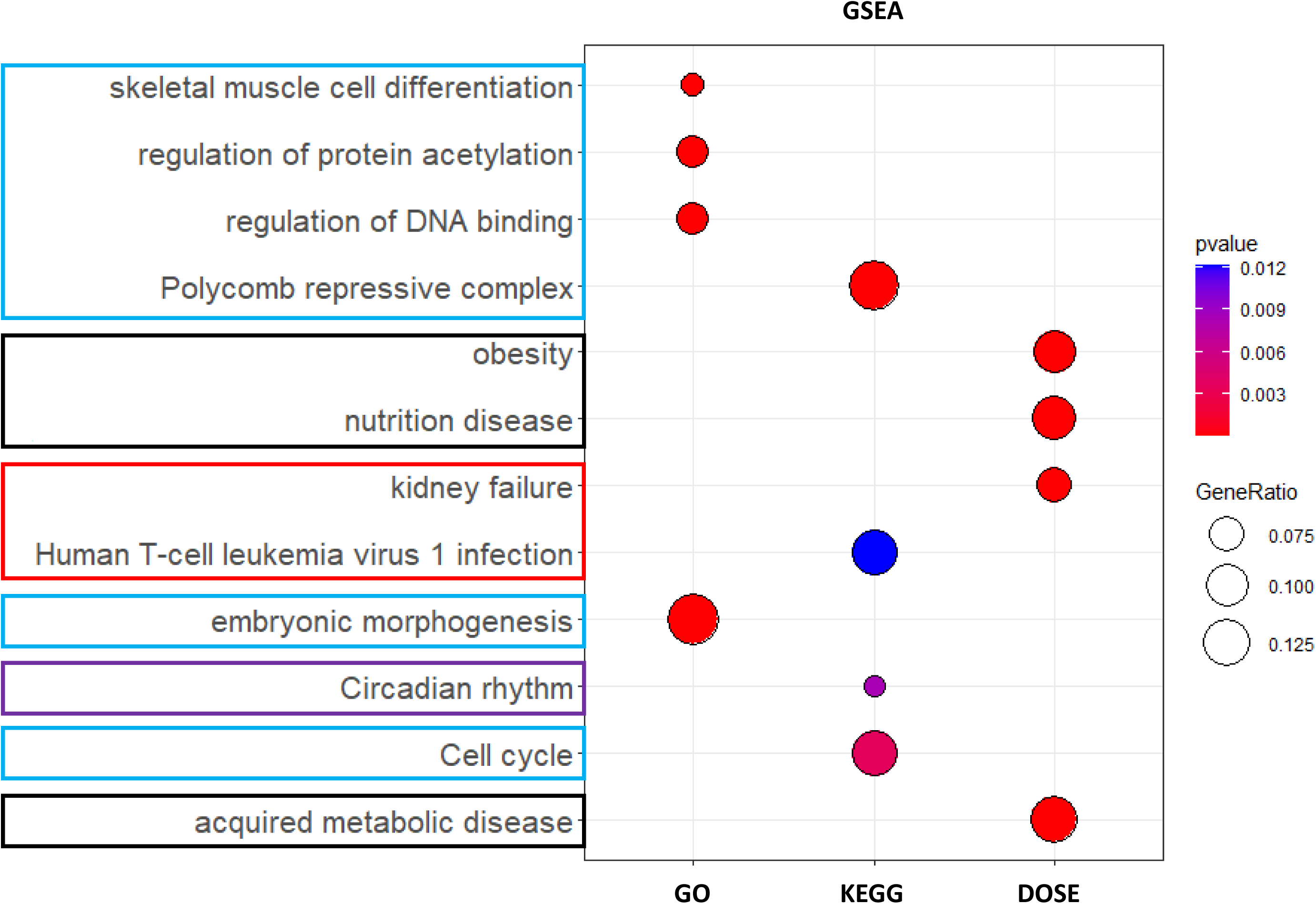
Uncovering Altered transcriptional regulation in HCC subtype of liver cancer through different Enrichment Analysis. Enrichment analysis were conducted to ascertain the communication pathways most significantly disrupted by HCC. This analysis involved comparing normal pathways to those that exhibited statistical differences within both the surviving and deceased populations, as well as between the groups of survivors and deceased individuals with statistically distinct gene expression patterns. This analysis shows the pathways that are enriched through three different libraries along the x-axis. The predicted transcription factors were found to affect the above pathways. Enrichment analyses were performed by employing three distinct enrichment libraries, aiming to pinpoint the HCC-specific TFs as prognostic biomarkers. Moving from left to right, we observe enriched analyses involving gene ontology (GO), Kyoto Encyclopedia of Genes and Genomes (KEGG), and Disease Ontology Semantic and Enrichment (DOSE). These analyses help identify the most significantly disrupted signal pathways in HCC, associated with these TFs. These analyses highlight pathways impacted by viral infections and inflammatory responses (indicated by red outlines), tumorigenesis (light blue outline), and hormonal responses (purple outlines). The color-emphasized pathways, which coincide with earlier GSEA results indicating these TFs as potential diagnostic and prognostic biomarkers, are believed to operate via transcriptional control mechanisms.

**Table 1.** The complete list of the prognostic and diagnostic biomarkers identified as Transcription Factors (TFs) in this study with our ML/DL algorism. Comprehensive list of 102 Transcription Factors (TFs) as Prognostic and Diagnostic Biomarkers Identified through an ML/DL-based SVM model, inclusive of All Statistical Significance.

| HCC ML Predicted TF |  |  |  |  |  |  |  |
| --- | --- | --- | --- | --- | --- | --- | --- |
| ABLIM3 | AFF3 | AGFG1 | AKIRIN1 | ANAPC15 | ARNTL2 | BACH2 | BATF |
| BCL10 | BCL3 | BCOR | BCORL1 | C11orf53 | C14orf180 | C16orf95 | C4orf36 |
| C5orf58 | CABYR | CARD19 | CASZ1 | CCDC71L | CDCA7L | CDCA8 | CDV3 |
| CRY2 | CSRNP1 | DCAF12L2 | DDX3X | DEPDC7 | DNAJC1 | DTWD1 | DUT |
| ELF4 | ETV3L | FAM217B | FBXO30 | FBXO31 | FOXO6 | GAS2L3 | GATA3 |
| GCHFR | HDAC2 | HJURP | HLF | HNF4A | HOXB13 | HOXC6 | HOXC8 |
| IL36B | KDM5D | KIAA0930 | KIAA1841 | KLF15 | KLF4 | LIMS2 | LMO4 |
| MACC1 | MAFF | MAML2 | MAP10 | MBNL3 | MCM10 | MED31 | MED8 |
| MPLKIP | MPND | MSANTD3 | MSC | MT-ATP8 | MYBL2 | MYC | MYCN |
| NFYB | NKX3-2 | NR0B1 | NR1H3 | NR6A1 | OCEL1 | ODF3L1 | OGFRL1 |
| PACRG | PAGR1 | POF1B | POLB | POLD4 | PPP1R10 | PPP1RC | RAVER2 |
| RFX2 | RING1 | RNF2 | RORC | RPS27L | S100A16 | SAMD5 | SAP30 |
| SERBP1 | SETDB2 | SFPQ | SOWAHC | SOX11 | SOX4 |  |  |

## Discussion

### Comparative analysis reveals divergent trends in for biomarkers for major subtypes of HCCs

Hepatocellular carcinoma (HCC) stands as a major contributor to cancer-related mortality and presents a significant global cancer-related concern, necessitating continuous efforts to enhance screening, diagnosis, and treatment [25–29,42]. Although alcohol- and nonalcoholic steatohepatitis (NASH)-related HCC burdens are rising, chronic viral hepatitis remains a global driver of HCC [25–29,42,74]. Viral-related HCC’s pathophysiology involves liver inflammation, oxidative stress, and disrupted cell signaling pathways. Viral oncogenicity stems from its DNA integration and persistence despite nucleotide analog suppression, highly relevant to transcriptional regulation [74–76]. Antiviral therapy reduces HCC risks, but advanced chronic liver disease patients require ongoing surveillance. Despite advancements in treatments, HCC remains highly lethal. Recent studies highlight immunotherapy’s potential in HCC treatment. Exosomes play a crucial role in negatively regulating the immune response in HCC, presenting new avenues for intervention.

### Pressing need for diagnostic and prognostic biomarkers specific to the HCCs

HCCs pose challenges due to their poor survival rates, particularly in advanced stages. Early detection significantly enhances survival chances through curative treatments. Current guidelines advise ultrasound (USG) and alpha-fetoprotein (AFP) monitoring for HCC screening in high-risk groups, while abdominal USG, magnetic resonance imaging (MRI), and magnetic resonance cholangiopancreatography (MRCP) are recommended for biliary tract cancer [74,76]. Nonetheless, advanced HCCs continue to develop in high-risk individuals despite these screenings. The quest for effective biomarkers for surveillance and early diagnosis of HCCs remains challenging [74,76–79]. Existing blood biomarkers, with limited sensitivity and variable specificity, are being developed to target high-risk subgroups of hepatocellular carcinomas (HCCs). Promising outcomes have emerged from studies using individual or combined markers like AFP, AFP-glycoform L3 (AFP-L3), des-gamma-carboxy prothrombin, glypican-3 (GPC3), osteopontin (OPN), midkine (MK), neopterin, squamous cell carcinoma antigen (SCCA), Mac-2-binding protein (M2BP), cyclic guanosine monophosphate (cGMP), and interleukin-6 (IL6) for HCC screening [77], thereby emphasizing the need for an expanded collection of potential HCC biomarkers. The current absence of dependable biomarkers for diagnosing and predicting hepatic cancers emphasizes the necessity to uncover new biomarkers for distinct subtypes, thereby enhancing opportunities for early detection and treatment for patients [11,12,15,16,19]. While AFP stands as the benchmark, it faces limitations in detecting varying grades and stages of HCC and accurately distinguishing between primary liver cancer types [15,17,80].

Biomarkers play a pivotal role in forecasting and managing liver cancer, facilitating early detection and foreseeing patient responses to treatment and disease progression [10,11]. Given liver cancer’s 5-year survival rates spanning 5% to 30%, timely detection significantly influences patient outcomes, particularly for HCC with survival rates are 15% [5]. Prompt HCC treatment correlates with extended periods free of recurrence and enhanced overall survival rates [13]. Nevertheless, chemotherapy resistance is linked to HCC, posing treatment challenges and leading to a grim prognosis [14]. Hence, pinpointing sensitive and specific biomarkers can aid in the timely diagnosis and control of liver cancer, ultimately ameliorating patient outcomes.

### Prognostic and diagnostic potentials of CmPn members for distinguishing major types of HCCs

Recently, we conducted a comprehensive analysis for the potential association between of the HCCs and the members of the CmPn network [20]. Leveraging two databases containing patient samples with differential expression data for these key CmPn components. Sociological, pathological, and follow-up clinical data for HCC patients extracted from the TCGA-LIHC and TCGA-PANCAN databases [30]. Our prior research outcomes unveiled strong clinical connections between all components of the CmPn network in HCC, while only a few members of CmPn network members with CCA [20]. Furthermore, our data unveiled a predominant down-regulation of CmPn genes in CCA, setting it apart from the up-regulation of CmPn genes in HCC [20]. This underscores the fundamental distinction between the two major liver cancer types, especially HCCs [20]. Moreover, rarer liver cancer subtypes have even scarcer options for early detection. Thus, our observation of varying expression of multiple components within the CmPn network in both HCC and CCA clinical tumor samples, at both gene expression and protein levels, underscores their potential as innovative biomarkers for liver cancer. These findings hold promise for creating more precise and sensitive biomarkers, aiding in early diagnosis, treatment response assessment, and prognosis for individuals with liver cancer. Lately, a growing body of evidence indicates that tumor cells can exhibit traits resembling those of stem cells. This imparts upon tumor cells the capability for self-renewal, which plays a role in the continual presence of tumors and encompasses factors like therapy resistance, dormancy, and the instigation of metastasis. These responses are triggered by diverse physiological cues, including hormone steroids [64,81–83].

In order to confirm our earlier discovery, we conducted a renewed evaluation of the complete constituents within the CmPn signaling network, utilizing an expanded dataset of HCC information and more sophisticated methodologies. Our findings demonstrated that all 14 elements of the CmPn pathway have the potential to function as prognostic indicators for HCC. Moreover, among these 14 elements, 9 exhibited the capability to serve as both prognostic and diagnostic markers (Fig. 2C), thus providing additional confirmation of our initial hypothesis. Our new validated data also align well with previous reports that progesterone receptor membrane component 1 (PGRMC1), a member of CmPn, is linked to HCC progression from tumor size G2 to G3 grades of HCC and explored as a prognostic biomarker [21,22]. Elevated CCM3 gene expression is associated with poor prognosis due to its role in cell proliferation and metastasis in HCC Altered expression of all three CCM (1-3) genes, along with PAQR7, in early HCC stages [20,23,24].

This study aimed to identify biomarkers through bioinformatics analysis of publicly available datasets. Using publicly accessible expression data from HCC patients, we merged gene expression and clinical information to construct Kaplan–Meier (KM) survival curves. These curves were generated to comprehend the impact of the CmPn network on the overall survival (OS) of HCC patients. In each illustration, the left KM curve originated from KMplotter [84], while the right counterpart was established through TCGA data to validate our Kmplotter analysis. We ultimately identified 647 candidate diagnostic and prognostic biomarkers associated with HCC patient age and survival. Pathway analysis helps elucidate functions and disrupted pathways correlated with these biomarkers. The study also explores the prediction of transcription factors (TFs) using a machine learning model called ESM-TFpredict [58,69], which lead to the identification of 102 potential TFs, leveraging protein sequence data. The dataset preprocessing involves curating TF sequences and non-TF sequences for analysis. The model’s experimental execution and evaluation demonstrate its reliability and precision in TF prediction. Overall, the study combines bioinformatics analysis, transcriptomic investigation, and machine learning to uncover potential diagnostic and prognostic biomarkers for HCC. The utilization of publicly available data enhances our understanding of liver cancer’s molecular profiles and clinical significance, particularly in the context of HCC.

Our evaluation disclosed that elevated expression of the indicated elements corresponded to significantly poorer prognostic outcomes in HCC patients. Among these six vital CmPn components, all survival curves were confirmed except for PAQR5, with validation conducted using TCGA data. These outcomes further affirm the involvement of the CmPn signaling network in liver cancer tumorigenesis, underscoring their potential utility as prognostic indicators in HCCs. Ultimately, these findings, coupled with survival curves from the TCGA database, lend further support to the prospective utilization of CmPn genes as prognostic biomarkers for individuals with HCC.

## Conclusion

The findings in this study are significant because early diagnosis and treatment can greatly improve HCC patient outcomes. The expanded pool of potential prognostic biomarkers, along with the identification of disrupted pathways and altered CmPn members in various HCC subtypes, offers a significant advantage. This abundance of candidates is invaluable for seeking HCC-specific grading and staging biomarkers, paving the way for targeted therapies to enhance patient outcomes and potentially improve overall survival rates. Moreover, these findings offer insights into liver cancer’s molecular mechanisms, potentially fostering an improved comprehension of the disease and innovative approaches for prevention and treatment.

## Ethics approval and consent to participate

Not applicable

## Consent for publication

Not applicable

## Author contributions

Detailed in the CREDIT author contribution information on the online submission system.

## Acknowledgement

We wish to thank Muaz Bhalli, Alexander Le, Ofek Belkin, Mellisa Renteria, David Jang, Justin Aickareth, Victoria Reid, Majd Hawwar, Revathi Gnanasekaran, Nickolas Sanchez, Charlie Harvey, and Drexell Vincent at Texas Tech University Health Science Center El Paso (TTUHSCEP) for their technical help during the experiments.

## Availability of data and materials

Some essential analytical data were provided in supplemental materials in the online version of the journal, while all data submitted were comply with Institutional or Ethical Review Board requirements and applicable government regulations.

