## Supplementary material for "Novel hepatocellular carcinomas (HCC) Subtype-Specific Biomarkers": Suppl Materials

Running title: HCC-Specific Biomarkers

**Supplemental Materials**

*All correspondence:*
Jun Zhang, Sc.D., Ph.D.

Department of Molecular & Translational Medicine (MTM)
Texas Tech University Health Science Center 5001 El Paso Drive, El Paso, TX 79905

**Supplemental Legend**

Supplemental Figure 1: **The analysis of greater than 19,000 protein coding RNA sequences were compared between the survivors and deceased populations to determine biomarker capacities**. The remaining Kaplan-Meier Survival curves of HCC Patients involved comparing these outcomes with noncancerous control biopsies to assess their potential as diagnostic biomarkers. **1A.** The remaining Kaplan-Meier Survival curves of HCC Patients for prognostic biomarkers, showing difference between the living and deceased HCC groups, and these biomarkers are an extension of those in figure 2A. **1B.** The remaining Kaplan-Meier Survival curves of HCC Patients for diagnostic and prognostic biomarker candidates exhibited statistically significant differences from both the survivor and deceased HCC cohorts, as well as the survivor HCC cohort and control populations, and the deceased and control populations, as an extension of those in figure 2B.

Supplemental Figure 2: **Over 900 biomarkers had significant prognostic candidacy, to explore how these processes affect the overall body system we used gene set enrichments to determine areas that are affected by hepatocellular carcinomas. 2A:** This gene set enrichment is from the Gene Ontogeny library the prognostic candidates were found to affect metabolic process, which was to be anticipated from the organ affected with cancer. Interestingly we begin to see wound healing and responses to wounds affected. **2B**: The next enrichment of these biomarker candidates was through the KEGG library and again we find the metabolism aspects of cellular process are affected. **2C:** The final enrichment library that was utilized in this analysis was through the disease ontogeny library. Many metabolic related processes have similar affected genes when compared to HCC.

Supplemental Figure 3: **Identification of 646 prognostic/diagnostic biomarkers for HCC and utilized gene set enrichments to see other related affected processes. 3A**: The first gene set enrichment utilizes the gene ontogeny library where we find a greater response to wounds and blood coagulation. This could provide insights on to how liver cirrhosis occurs with HCC. **3B:** We utilized multiple library enrichments for this analysis, the next part of this analysis occurred after utilizing the KEGG library. This enrichment seems to have more of a response to hormone then in the previous biomarker sets. **3C:** The final library was the disease ontogeny library where nutrition and metabolic process are still affected.

Supplemental Figure 4: **We utilized machine learning to identify possible 102 transcription factors (TFs) from the 646 prognostic/diagnostic biomarkers**. **4A**: This gene set enrichment utilizes the gene ontogeny library where we found multiple pathways affected at the transcription level, which coincides with what we utilized machine learning to isolate. **4B**: Next, we utilized the KEGG library for gene set enrichments. Interestingly we begin to see inflammatory bowel disease as this is part of when patients become aware of symptoms associated with HCC. **4C**: Finally, we utilized a final gene set library of disease ontogeny. Interestingly we find many of the disease associated with the transcription factors enriched are found in the male reproductive and urinary system.

Supplemental Table 1: A comprehensive list of all prognostic and diagnostic biomarkers that was discovered during out analysis of the patient transcriptome.

Supplemental Figure 1A: Significant Prognostic Biomarkers for HCC

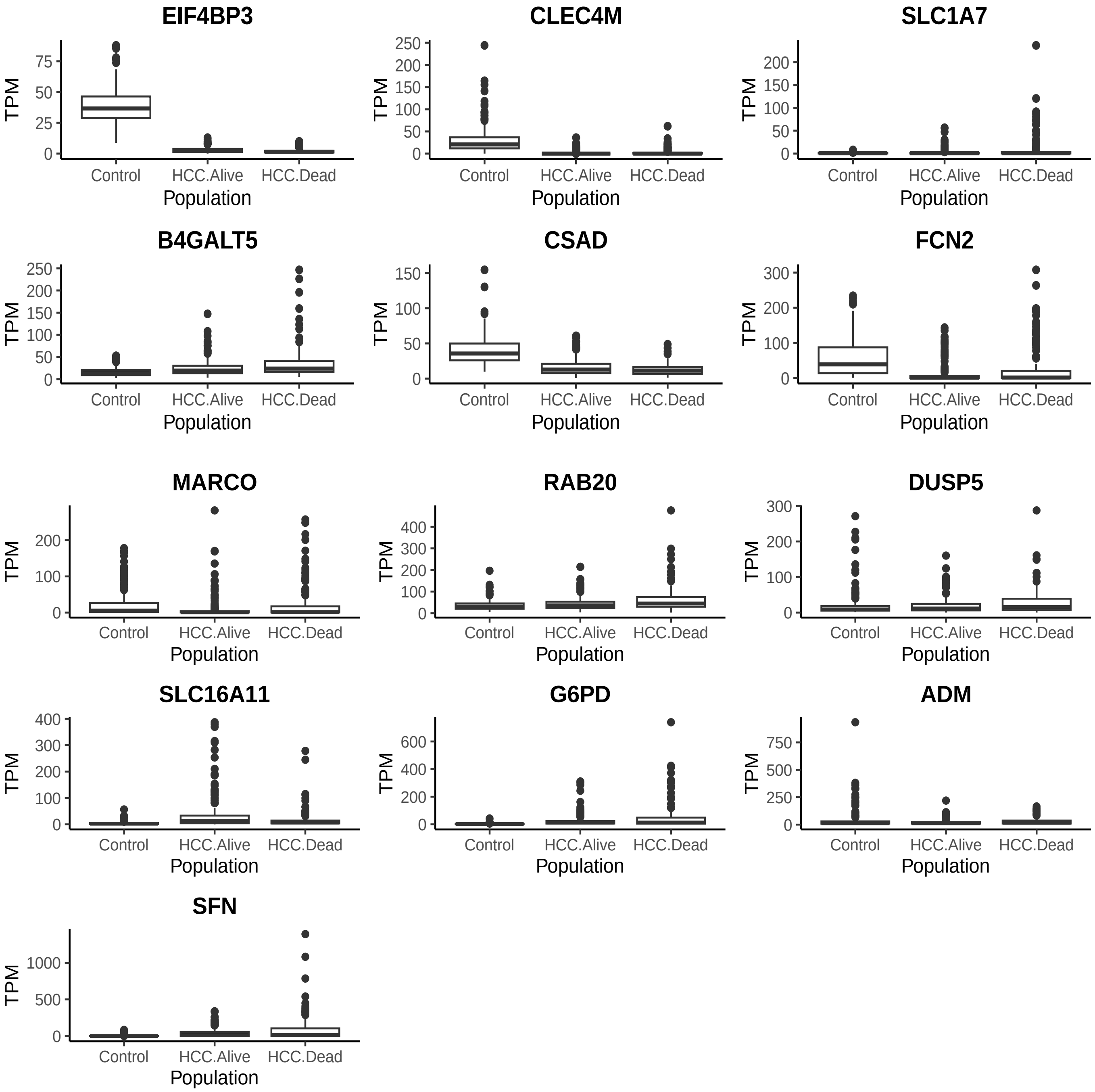

Supplemental Figure 1B: Significant Prognostic and Diagnostic Biomarkers for HCC

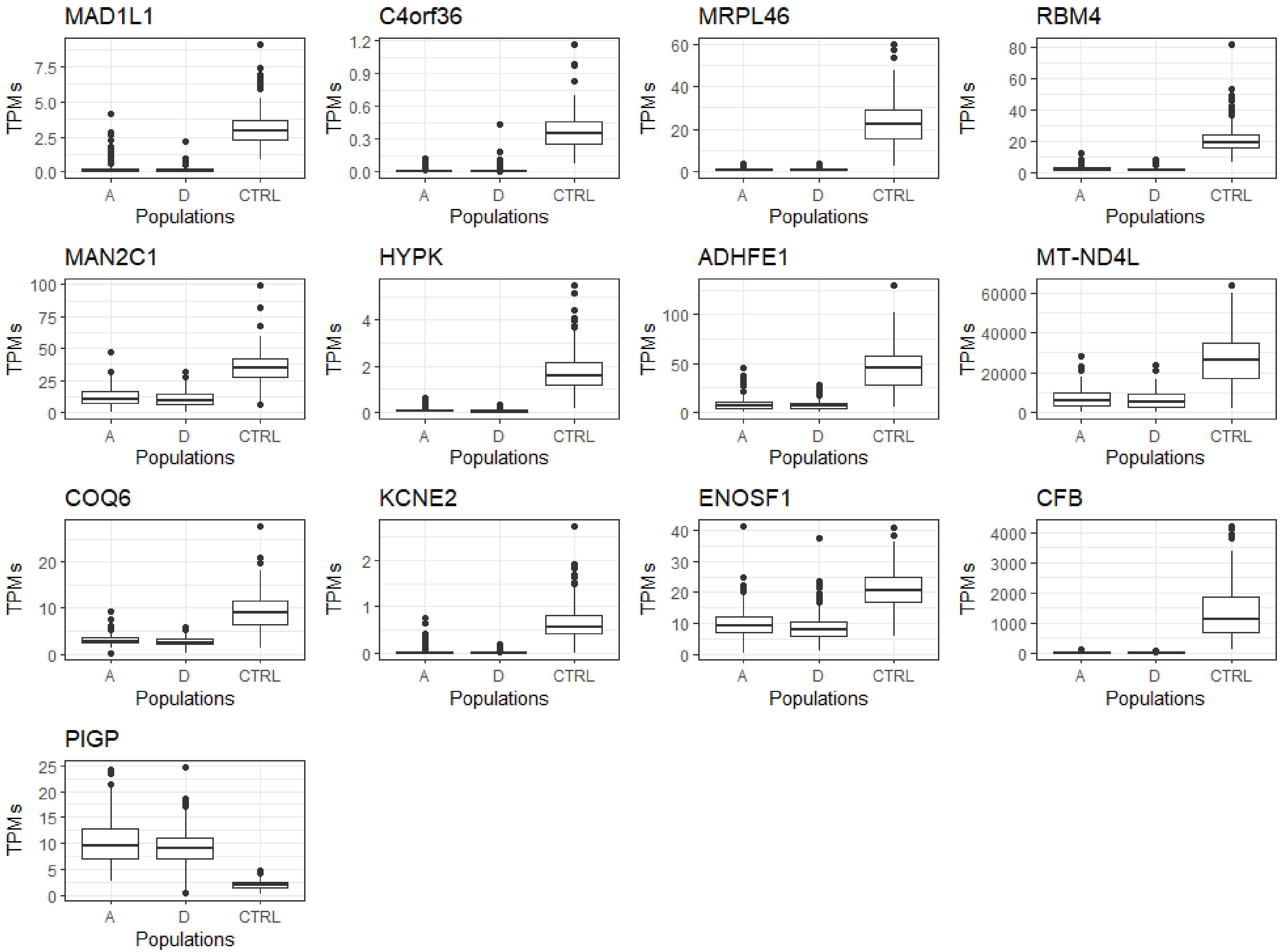

Supplemental Figure 2A: EnrichGO Prognostic only biomarkers

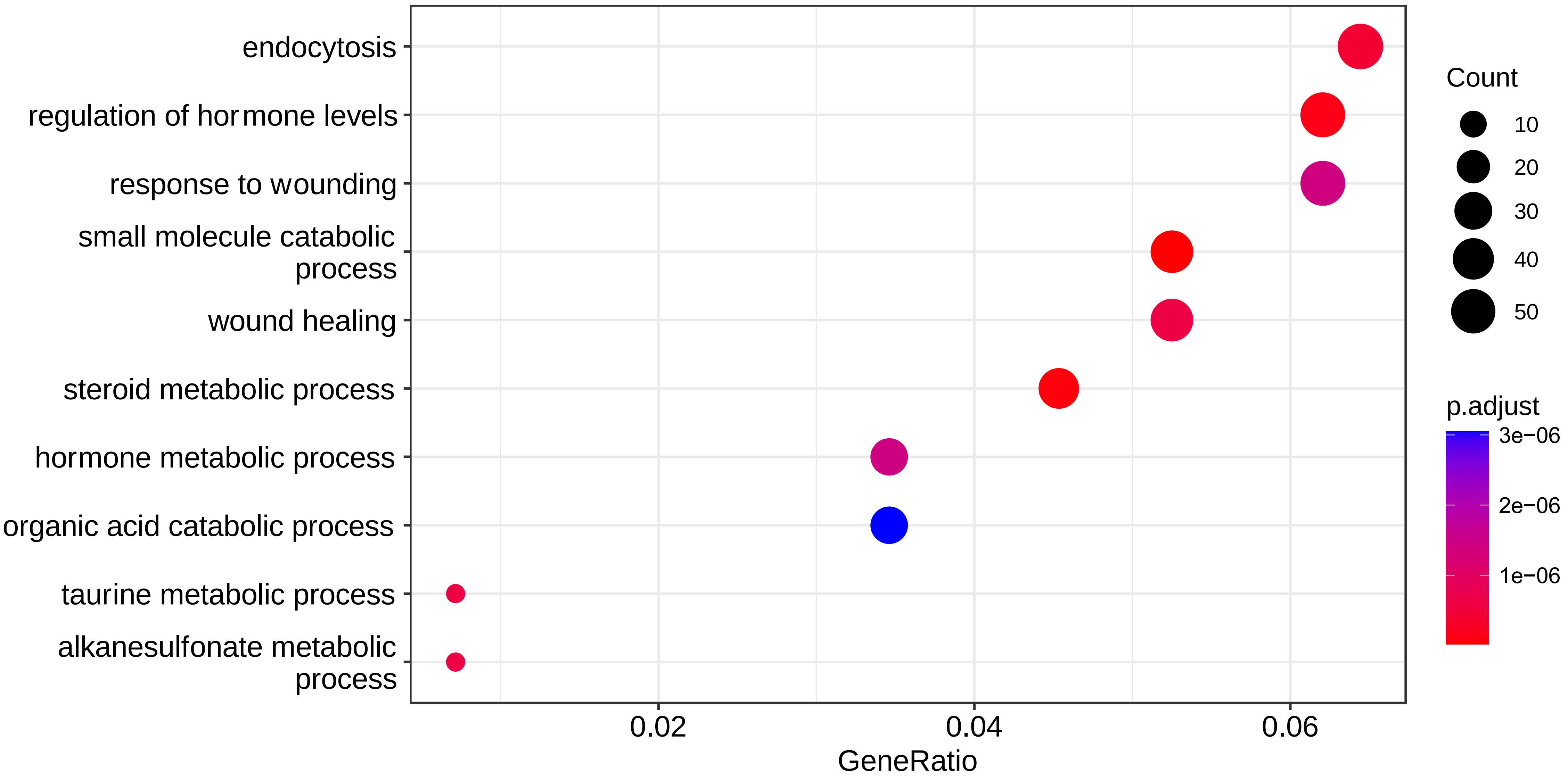

Supplemental Figure 2B: EnrichKEGG Prognostic only biomarkers

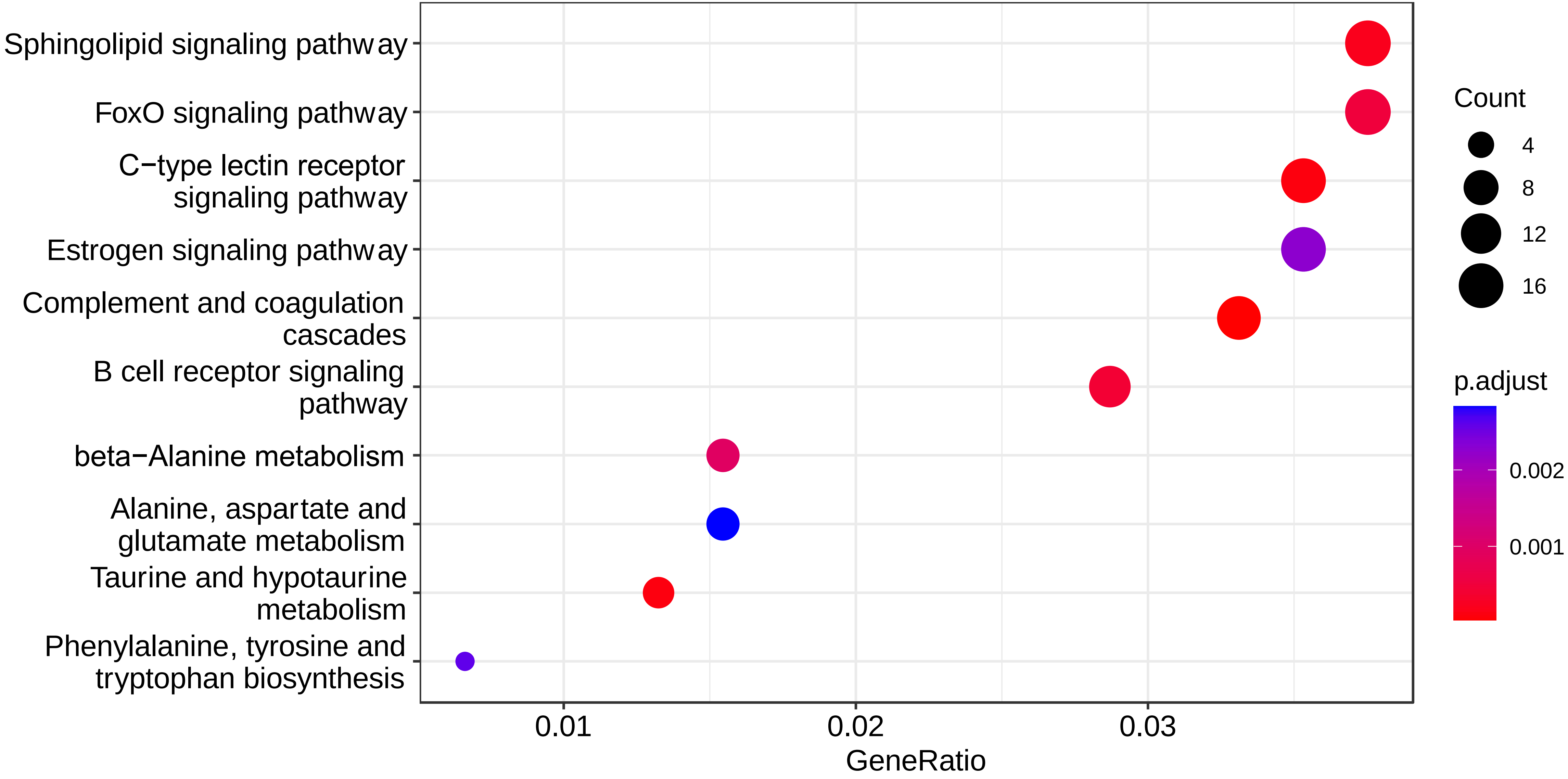

Supplemental Figure 2C: EnrichDO Prognostic only biomarkers

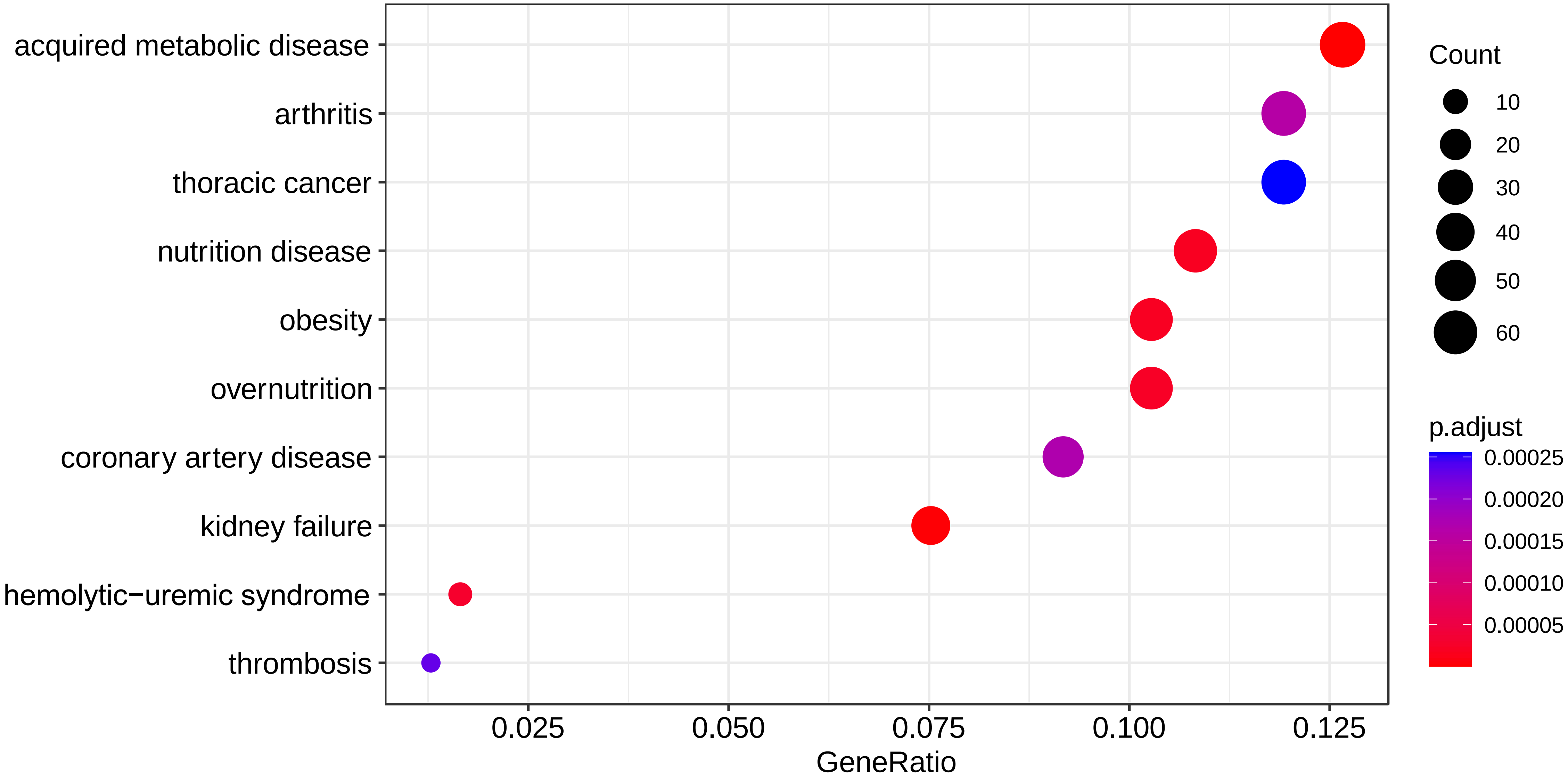

Supplemental Figure 3 A:
Diagnostic/Prognostic GO

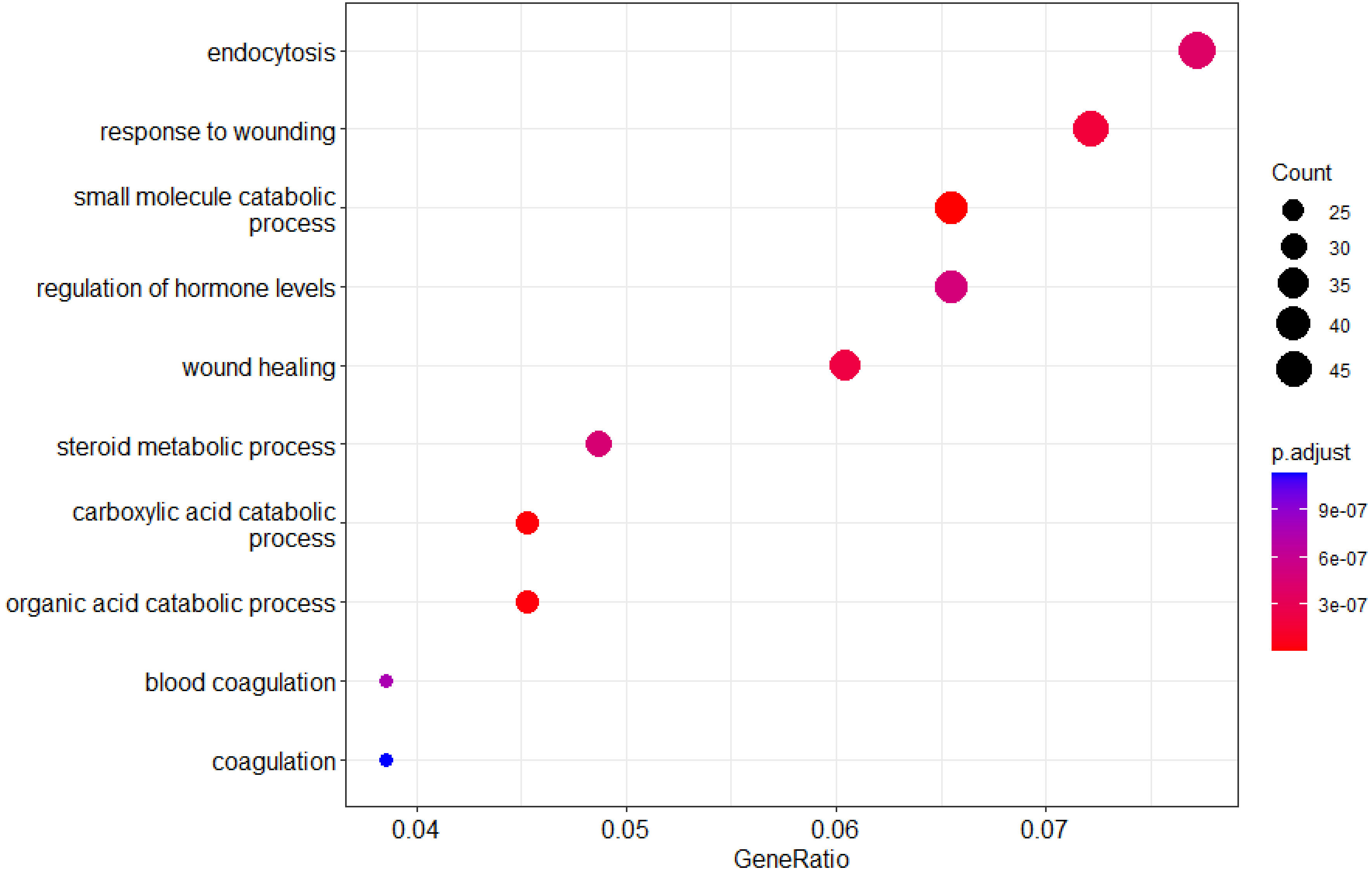

Supplemental Figure 3B:
Diagnostic/Prognostic KEGG

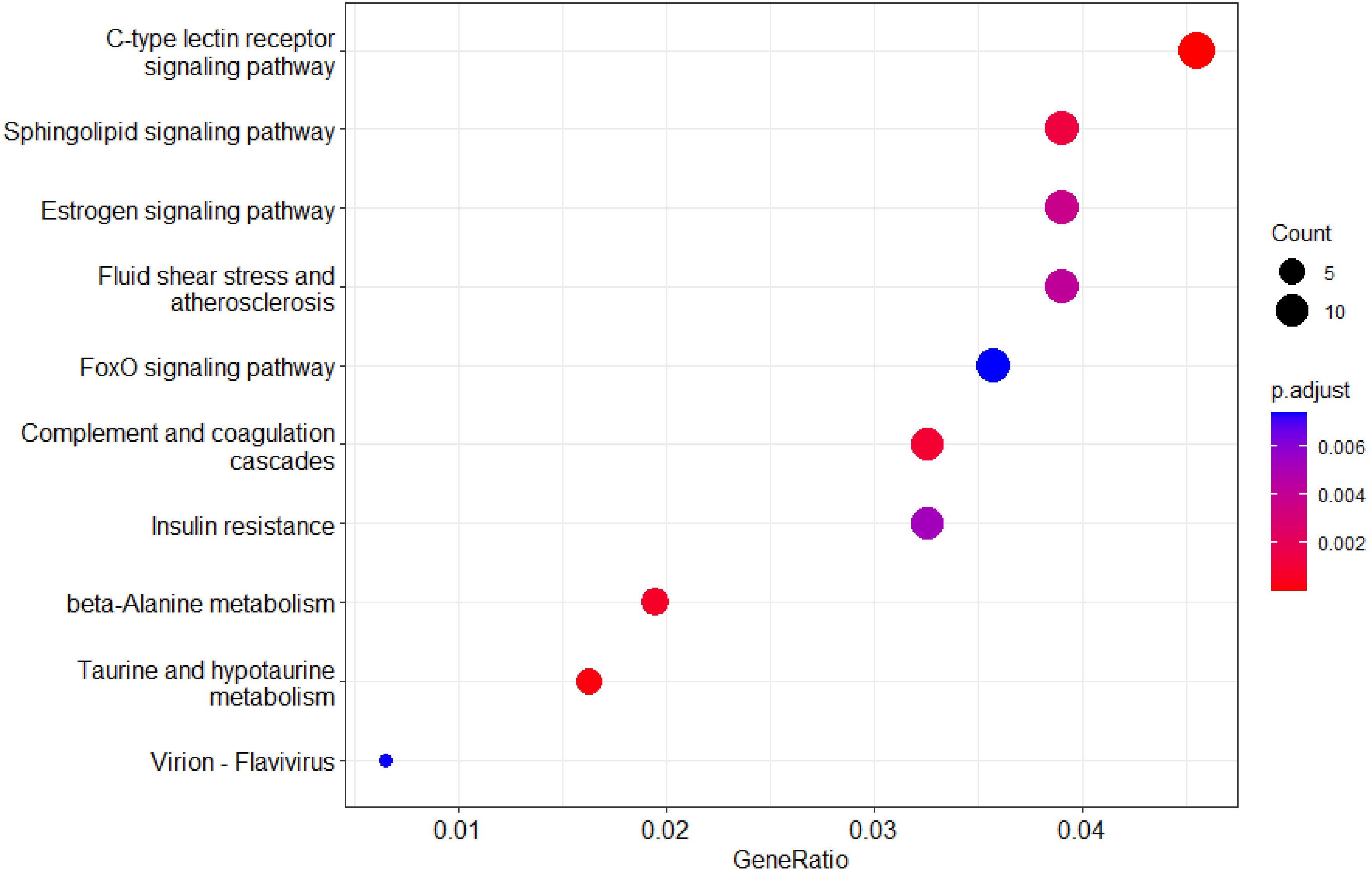

Supplemental Figure 3C:
Diagnostic/Prognostic DOSE

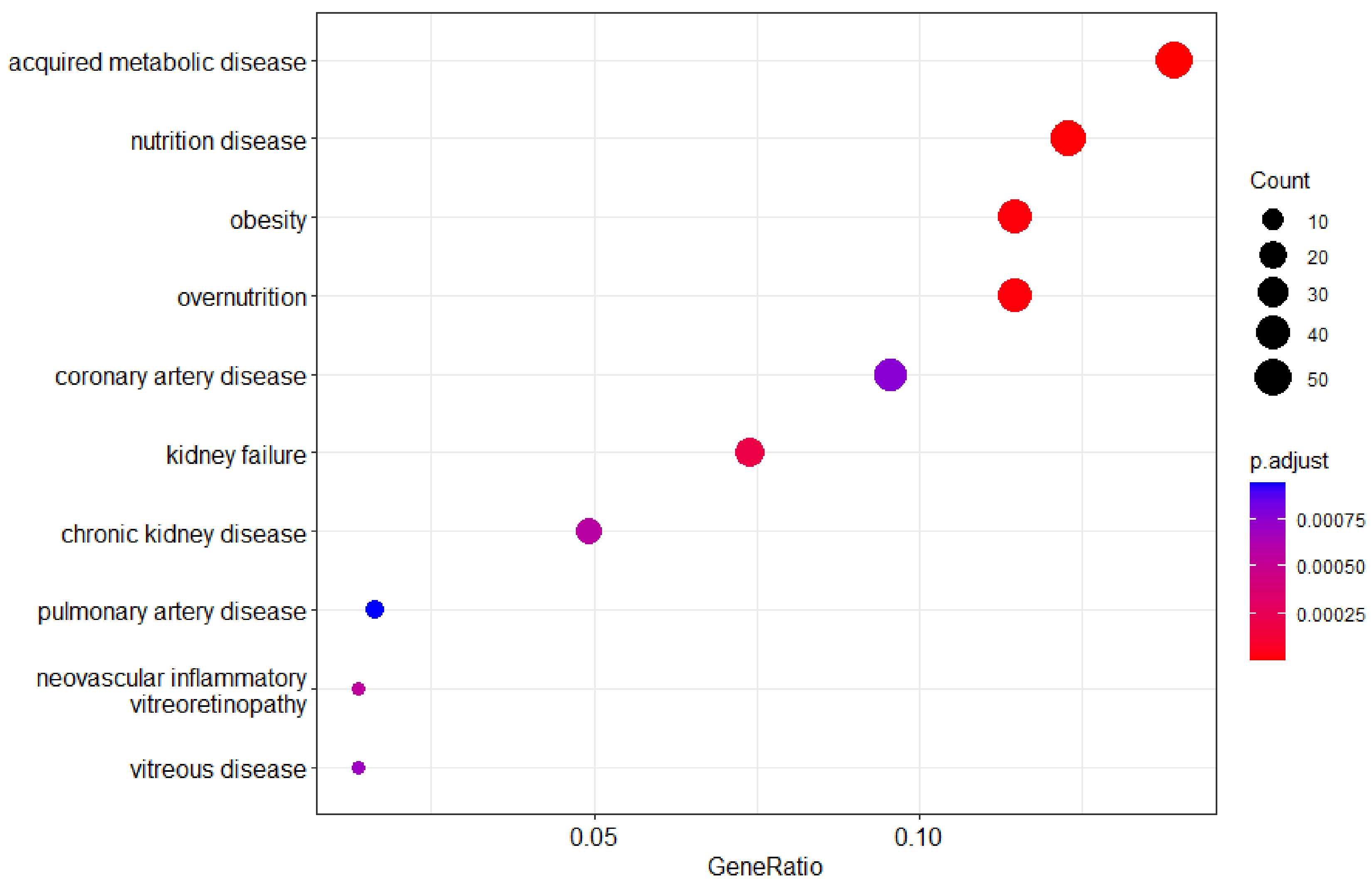

Supplemental Figure 4A: ML predicted TF of the prognostic and diagnostic biomarkers EnrichGO

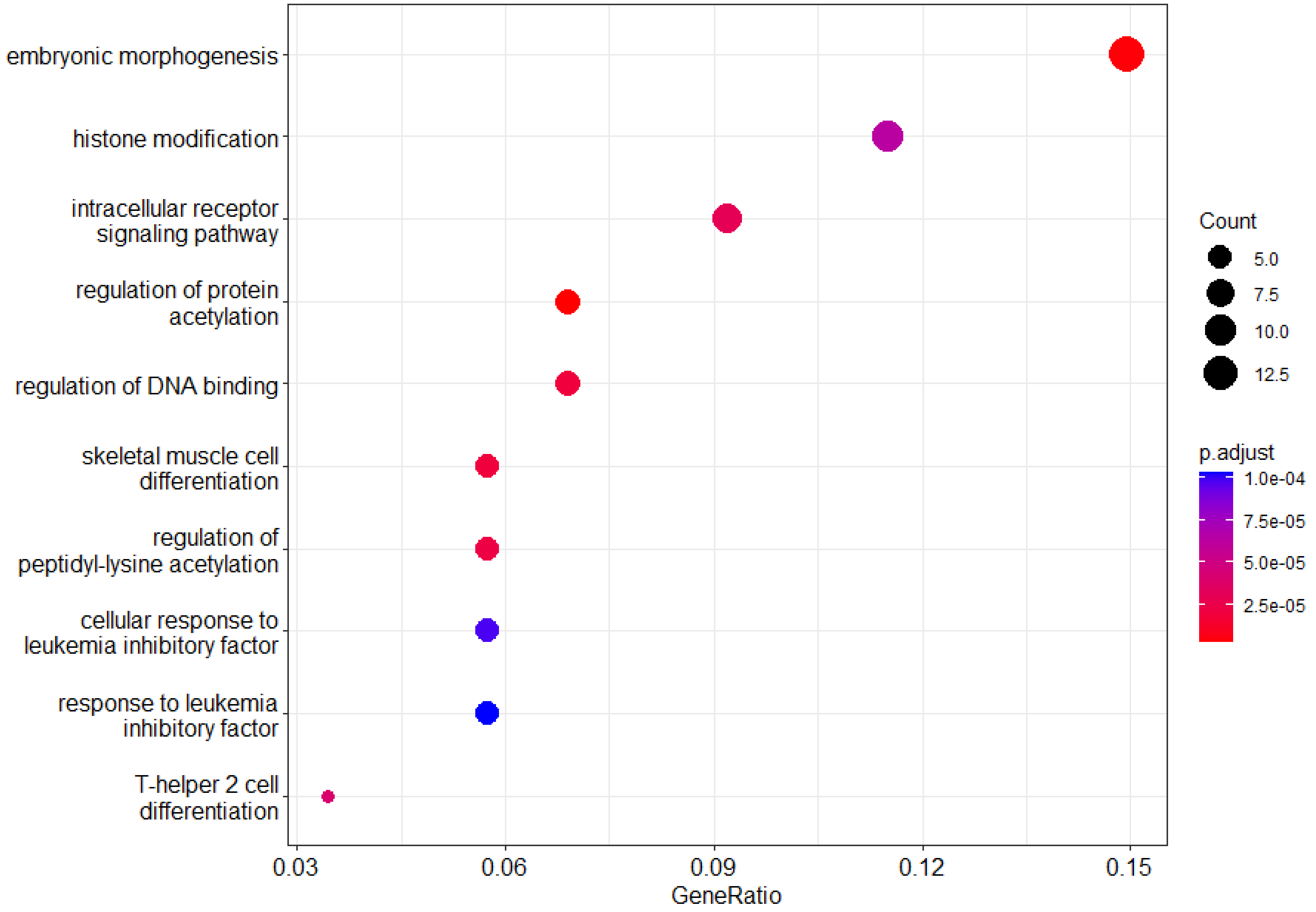

Supplemental Figure 4B: ML predicted TF of the prognostic and diagnostic biomarkers EnrichKEGG

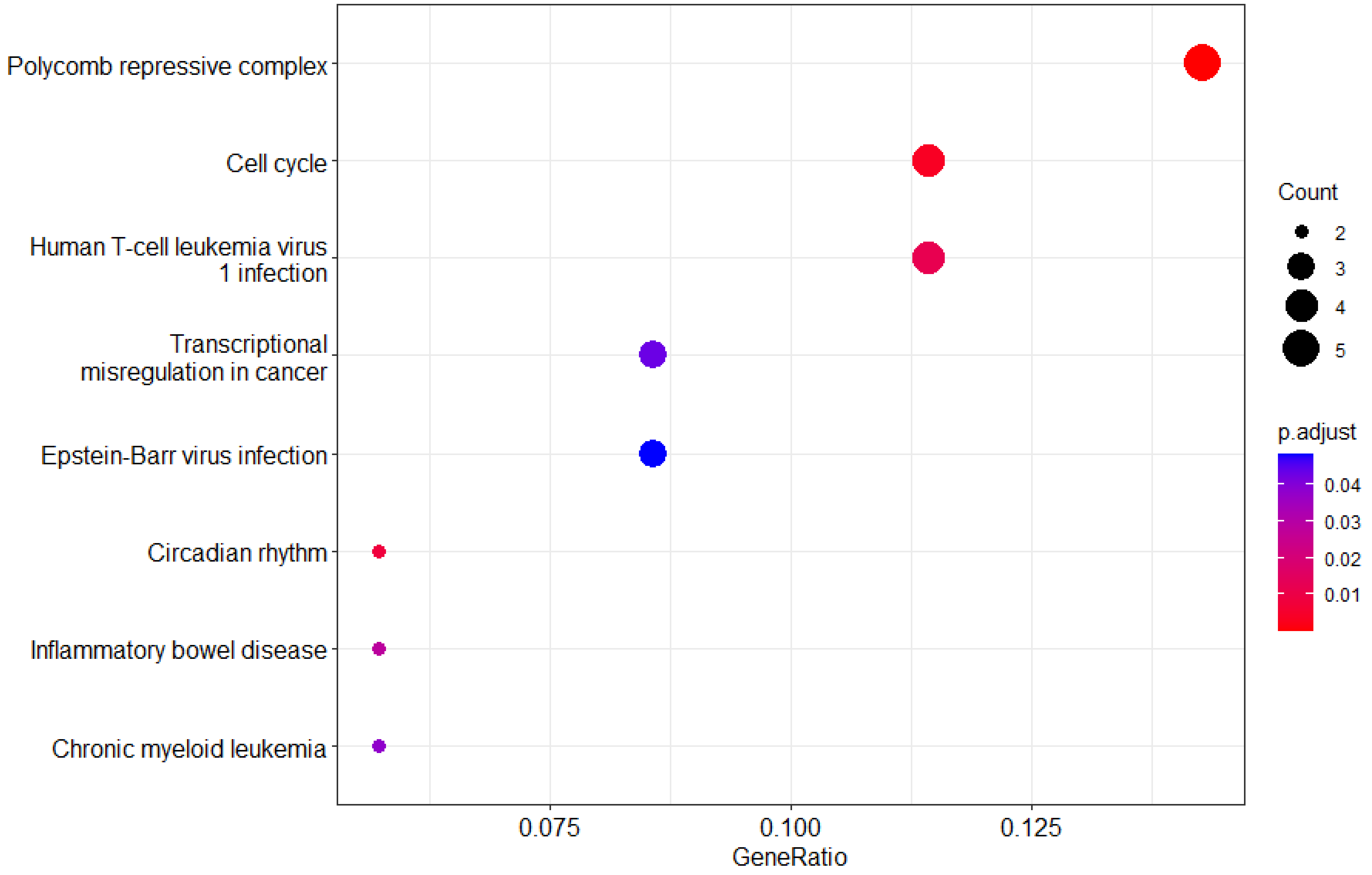

Supplemental Figure 4C: ML predicted TF of the prognostic and diagnostic biomarkers EnrichDO

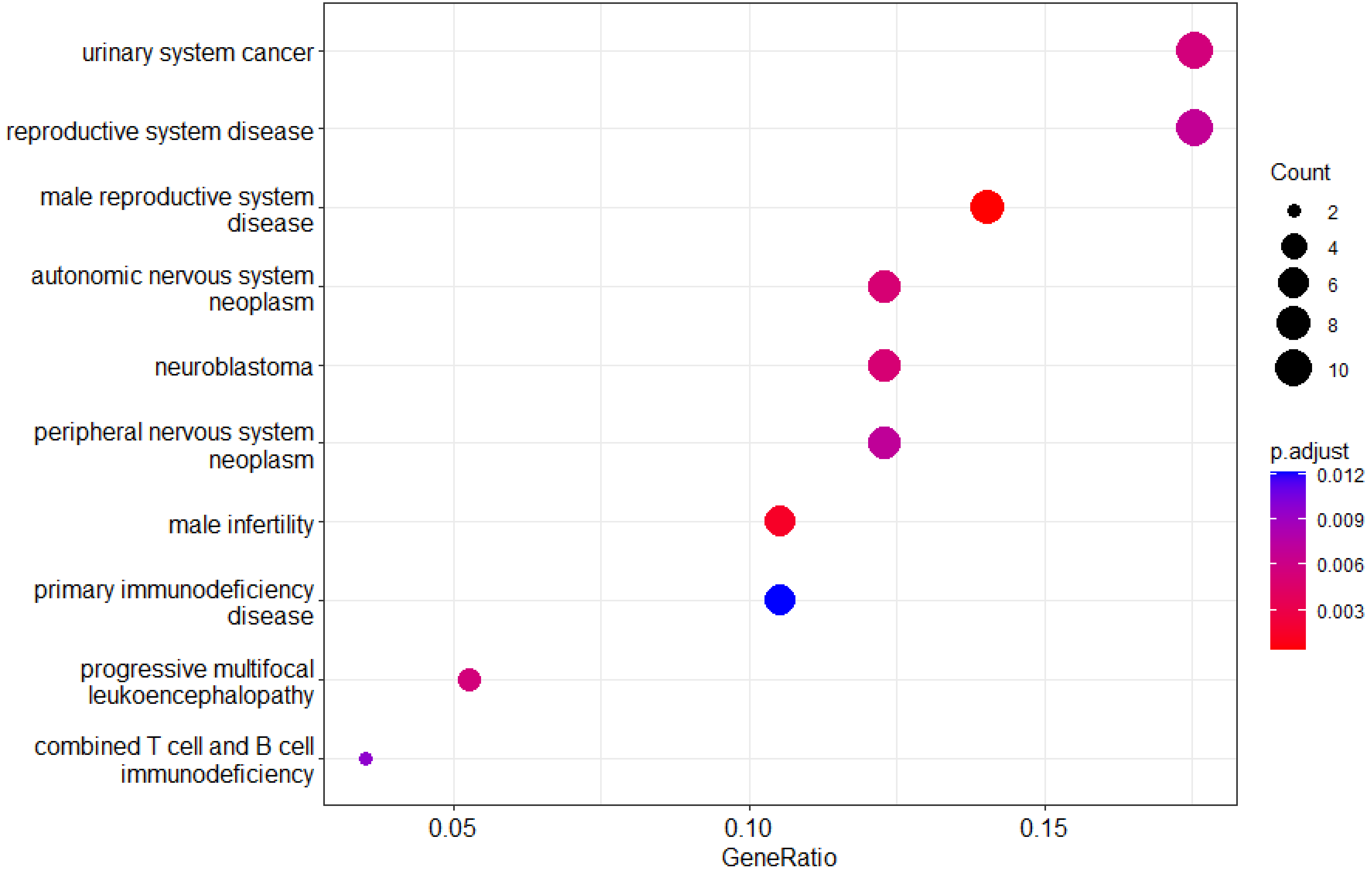

### Supplemental Table 1: Complete list of all HCC Prognostic/Diagnostic Biomarkers (Count:646)

| HCC Prognostic and Diagnostic Biomarkers | | | | | | | | | | | | | | | | | | | | | | | | | |
|---|---|---|---|---|---|---|---|---|---|---|---|---|---|---|---|---|---|---|---|---|---|---|---|---|---|
| AADAT | ALDH7A1 | B3GNT9 | C4BPB | CCDC43 | CEP97 | COX17 | DEPDC7 | EDN1 | FAM161A | FSD1L | GNPDA1 | HMCN2 | IQCD | KNG1 | MACC1 | MPLKIP | MYCN | NR1H3 | PCSK5 | PPFIBP2 | PZP | RIT1 | SCRN1 | SHISA9 | SLC39A10 |
| ABCC1 | ALG5 | B4GALT5 | C4orf36 | CCDC71L | CERK | COX5A | DGKG | EFNB1 | FAM180A | FUNDC2 | GPD2 | HMGCS1 | IQSEC1 | KPNA2 | MAD1L1 | MPND | MYO18B | NR6A1 | PEBP1 | PPIB | RAB20 | RMDN1 | SCRN2 | SIAH1 | SLC44A1 |
| ABCC5 | ANAPC15 | BAAT | C5AR1 | CCL15 | CFB | CPB2 | DGKQ | EIF4EBP3 | FAM20A | FZD7 | GPR35 | HMMR | IRAK3 | KRT17 | MAFF | MPP6 | N4BP3 | NRAS | PECR | PPIL3 | RAB32 | RNF19B | SDC2 | SIAH2 | SLC48A1 |
| ABLIM3 | ANGPTL6 | BACH2 | C5orf58 | CCL16 | CFHR1 | CRHBP | DHDH | ELF4 | FAM217B | G6PC | GPR84 | HNF4A | IRS1 | KRT81 | MAGEB6 | MPV17 | NABP1 | NRG1 | PET100 | PPM1G | RAB3IL1 | RNF2 | SDC4 | SIGLEC7 | SLC6A1 |
| ACOT13 | ANKRD13B | BAIAP2 | CA14 | CCM2L | CFP | CRY2 | DHRS12 | ELOVL2 | FAM3B | G6PC3 | GPRC5B | HOMER1 | ISG20L2 | KRTAP11.1 | MAML2 | MPZL1 | NARF | NSUN4 | PEX13 | PPP1CB | RAB4A | ROR2 | SEC14L2 | SIK2 | SLC7A7 |
| ACOXL | ANKRD24 | BATF | CABYR | CCR1 | CGNL1 | CSAD | DIO2 | EMC4 | FAM8A1 | G6PD | GPRIN1 | HOXB13 | ITGAM | KRTAP4.1 | MAN1A1 | MRPL37 | NBPF6 | NTF3 | PEX5 | PPP1R10 | RAB6B | RORC | SELENOI | SKAP2 | SLC9A3R1 |
| ACTG1 | ANLN | BCL10 | CADM1 | CCT6B | CHAC2 | CSF3R | DISP1 | EMC6 | FBXL2 | GAD1 | GPSM2 | HOXC6 | ITGB7 | KRTAP4.12 | MAN2C1 | MRPL46 | NCF2 | NXF3 | PFKFB4 | PPP1R1C | RAB7A | RP2 | SEMA6A | SLC11A1 | SLC9C1 |
| ADAMTS13 | ANOS1 | BCL3 | CALM1 | CD24 | CHIC2 | CSRNP1 | DLGAP3 | EMCN | FBXL5 | GADD45B | GRK3 | HOXC8 | ITIH1 | LDAH | MAP10 | MRPL9 | NCOA7 | OCEL1 | PGD | PPP2R1B | RAB7B | RPL30 | SEMA7A | SLC16A11 | SLCO2A1 |
| ADAMTS5 | ANP32B | BCO2 | CALML3 | CD300A | CHRM2 | CSTF2 | DNAJC1 | ENDOG | FBXL7 | GALR3 | GRPEL2 | HP | ITPR2 | LGR5 | MAP1LC3B2 | MRPS18C | NDC1 | OCIAD1 | PHKA2 | PPP2R3A | RAC1 | RPS27L | SEPHS1 | SLC19A3 | SMIM10 |
| ADAP2 | AP1S3 | BCOR | CAMK2N1 | CD300C | CHST9 | CTSL | DNAJC5G | ENOSF1 | FBXO30 | GAS2L3 | GSAP | HS6ST1 | JMY | LHFPL2 | MAP7D3 | MRPS28 | NDRG3 | ODAM | PIGP | PPT1 | RASL10B | RPUSD2 | SERBP1 | SLC1A7 | SMPD1 |
| ADGRE5 | APOE | BCORL1 | CAMSAP1 | CD5L | CILP2 | CTTNBP2NL | DNASE1L3 | ENTPD2 | FBXO31 | GATA3 | GSR | HSD17B4 | KCNE2 | LIFR | MARCKSL1 | MRPS31 | NDUFA4 | ODF3L1 | PIGS | PRDX1 | RASSF8 | RRP8 | SERF1A | SLC22A25 | SMS |
| ADHFE1 | APOL5 | BDH1 | CAP2 | CDADC1 | CLCF1 | CX3CL1 | DPYSL4 | EPHX1 | FBXO5 | GCHFR | GTPBP4 | HSD17B7 | KCNJ5 | LILRA6 | MASTL | MSANTD3 | NDUFAF1 | OGFRL1 | PIK3CB | PRELID2 | RAVER2 | RTN3 | SERPINB8 | SLC23A2 | SNAP25 |
| AFF3 | AQP6 | BMP10 | CAPN11 | CDC20 | CLDN2 | CXCL12 | DSTN | EPO | FCER1G | GCLM | HACD3 | HSD17B8 | KCNK1 | LILRB4 | MBNL3 | MSC | NDUFC1 | OPN1SW | PLA2G7 | PRKCD | RBM17 | RTP3 | SERPINB9 | SLC25A13 | SNX7 |
| AGFG1 | AQP9 | BMPER | CARD19 | CDC25B | CLDN4 | CYB5R2 | DTNBP1 | EPS8L3 | FCGR2B | GCNT4 | HADHA | HSPA12A | KDM5D | LIMS2 | MCC | MSR1 | NDUFC2 | OR8A1 | PLB1 | PROC | RBM4 | RXFP1 | SERPINC1 | SLC25A15 | SORD |
| AGTRAP | ARHGEF35 | BRSK1 | CARMIL1 | CDCA3 | CLEC1B | CYP11A1 | DTWD1 | ERI1 | FCGR3A | GDF2 | HAGH | HTRA3 | KIAA0930 | LMO4 | MCCC2 | MT.ATP8 | NECTIN1 | ORM2 | PLIN2 | PRPF38A | RBM47 | S100A10 | SETDB2 | SLC25A2 | SOWAHC |
| AKAIN1 | ARL15 | BRSK2 | CARS2 | CDCA7L | CLEC4D | CYP26A1 | DUSP15 | ESD | FCN2 | GDI2 | HAMP | HYPK | KIAA1217 | LPCAT1 | MCEE | MT.ND4L | NECTIN3 | OTUD1 | PLPPR1 | PRR11 | RDH8 | S100A11 | SF3B4 | SLC25A24 | SOX11 |
| AKIRIN1 | ARMCX1 | BSG | CASKIN1 | CDCA8 | CLEC4M | CYP26B1 | DUSP6 | ETV3L | FCN3 | GINS1 | HAVCR1 | IFITM2 | KIAA1841 | LRP10 | MCM10 | MTFMT | NEK3 | OXTR | PLSCR4 | PSD4 | RFPL1 | S100A12 | SFN | SLC25A42 | SOX4 |
| AKR1B15 | ARNTL2 | C11orf53 | CASP7 | CDKN1A | CMTM7 | CYP27A1 | DUT | EVA1A | FFAR4 | GIT1 | HBEGF | IFT172 | KIAA2013 | LRP12 | ME1 | MTHFD2L | NET1 | PACRG | PLXNA3 | PSRC1 | RFX2 | S100A16 | SFPQ | SLC25A47 | SPA17 |
| AKR1C1 | ASAH2 | C14orf180 | CASZ1 | CDO1 | CNDP1 | CYP2C19 | DYNLT1 | EXOC3L2 | FLNA | GLP1R | HDAC2 | IGFALS | KIF17 | LRP5 | MED31 | MTHFS | NFYB | PAGR1 | PNMA6A | PTCHD4 | RGCC | S100A6 | SFT2D1 | SLC2A2 | SPARCL1 |
| AKR1E2 | ASAP1 | C15orf40 | CAV2 | CDV3 | CNEP1R1 | CYP3A5 | EBAG9 | EZR | FMN2 | GLP2R | HDDC3 | IGSF3 | KIF18B | LRRC40 | MED8 | MTMR2 | NKAIN1 | PALM | PNP | PTDSS2 | RHBDL2 | S1PR1 | SFTPD | SLC2A4 | SPP1 |
| AKT2 | ASB4 | C15orf61 | CBLN1 | CEBPZOS | COG8 | DAB2 | EBPL | F13B | FMO3 | GLT8D1 | HILPDA | IL15RA | KIF20A | LRRC41 | MFSD2B | MTMR9 | NKX3.2 | PARD6B | POF1B | PTER | RHBG | SAMD15 | SFXN3 | SLC30A10 | SPPL2A |
| ALAD | ATG101 | C16orf95 | CBS | CENPI | COLEC10 | DCAF12L2 | ECI1 | F5 | FMO4 | GLUL | HINT2 | IL17D | KIF2C | LRRD1 | MGST2 | MTRF1L | NOL10 | PARP15 | POLA1 | PTGFRN | RHO | SAMD5 | SGCB | SLC35F6 | |
| ALAS1 | AXL | C1orf50 | CCDC110 | CENPO | COLGALT2 | DCAF8L1 | ECI2 | F8A3 | FNDC4 | GNAI3 | HJURP | IL20RA | KLF15 | LY6H | MIA3 | MYADM | NPTX2 | PARP16 | POLB | PTGS1 | RHOC | SAP30 | SGPP2 | SLC35G2 | |
| ALDH1B1 | B3GAT3 | C1orf53 | CCDC125 | CEP126 | COQ6 | DDX3X | ECM1 | FABP6 | FOXO6 | GNG4 | HK3 | IL36B | KLF4 | LYPLAL1 | MOB3B | MYBL2 | NQO1 | PCDH1 | POLD4 | PTH1R | RIMKLA | SBK3 | SGSM1 | SLC37A1 | |
| ALDH5A1 | B3GNT7 | C3orf52 | CCDC170 | CEP131 | CORO1C | DENND6A | ECM2 | FAM124A | FRRS1L | GNG7 | HLF | IMMP2L | KLRB1 | LYRM9 | MOCS2 | MYC | NR0B1 | PCDHGB2 | PON1 | PTP4A2 | RING1 | SCIN | SH3BP4 | SLC38A3 | |
